# The cytoskeleton as a smart composite material: A unified pathway linking microtubules, myosin-II filaments and integrin adhesions

**DOI:** 10.1101/195495

**Authors:** Nisha Bte Mohd Rafiq, Yukako Nishimura, Sergey V. Plotnikov, Visalatchi Thiagarajan, Zhen Zhang, Meenubharathi Natarajan, Shidong Shi, Virgile Viasnoff, Gareth E. Jones, Pakorn Kanchanawong, Alexander D. Bershadsky

## Abstract

The interrelationship between microtubules and the actin cytoskeleton in mechanoregulation of integrin-mediated adhesions is poorly understood. Here, we show that the effects of microtubules on two major types of cell-matrix adhesions, focal adhesions and podosomes, are mediated by KANK family proteins connecting the adhesion protein talin with microtubule tips. Both total microtubule disruption and microtubule uncoupling from adhesions by manipulations with KANKs trigger a massive assembly of myosin-IIA filaments. Myosin-IIA filaments, augmenting the focal adhesions and disrupting the podosomes, are indispensable effectors in the microtubule-dependent regulation of integrin-mediated adhesions. Myosin-IIA filament assembly depends on Rho activation by the RhoGEF, GEF-H1, which is trapped by microtubules when they are connected with integrin-mediated adhesions via KANK proteins but released after their disconnection. Thus, microtubule capturing by integrin-mediated adhesions modulates the GEF-H1-dependent effect of microtubules on the myosin-IIA filaments. Subsequent actomyosin reorganization then remodels the focal adhesions and podosomes, closing the regulatory loop.

## Introduction

The cytoskeleton is a composite material that is comprised of several types of fibrillar structures and molecular motors ^1^. The coordinated function of these cytoskeletal elements is poorly understood. We focus here on the crosstalk between microtubules and the actin cytoskeleton underlying many aspects of cell morphogenesis and migration. In particular, cell-matrix adhesions that can be considered as peripheral domains of the actin cytoskeleton are well known to be controlled by the microtubule system in many cellular models. Microtubule-driven alterations in cell-matrix adhesions are thought to be responsible for many microtubule-mediated changes in cell shape, polarization and migration ^2-7^.

Focal adhesions are elongated micron-sized clusters of transmembrane integrin molecules associated with a complex network of so-called plaque proteins that connect them to bundles of actin filaments ^8-11^. Mature focal adhesions evolve from small nascent adhesions that form underneath the lamellipodia in a force-independent manner. However, for maturation into focal adhesions, application of mechanical force generated by actomyosin contractility and formation of actin bundles transmitting such forces are required ^12-17^. It was shown long ago that disruption of microtubules by microtubule-depolymerizing agents leads to a substantial increase in the size and in some cases the number of mature focal adhesions ^18,19^, whilst microtubule outgrowth following washout of microtubule-depolymerizing drugs leads to a transient decrease in size or even the disappearance of focal adhesions ^20^. In addition, growing microtubules are targeted to focal adhesions so that their plus ends form multiple transient contacts with these structures ^21,22^. Several mechanisms for such targeting have been suggested ^5,15,23-25^. In particular, recent studies revealed that a novel component of focal adhesions, talin-binding proteins KANK1 and KANK2, can physically connect focal adhesions and microtubule tips via interaction with microtubule plus end-tracking proteins ^26-28^.

Podosomes represent another type of cell-matrix adhesion common in cells of monocytic origin but found also in cells of several other types ^29-31^. The characteristic structural feature of a podosome is an actin core perpendicular to the ventral cell membrane formed due to Arp2/3-dependent actin polymerization. The actin core is associated with an approximately ring-shaped cluster of adhesion proteins including transmembrane integrin molecules and the plaque molecules (paxillin, talin, vinculin etc.) linking integrins to the actin network, essentially similar to those in focal adhesions. In recent studies, myosin-IIA filaments were detected at the rim of podosomes and were claimed to participate in this link ^32-34^. Importantly, in contrast to the case for focal adhesions, disruption of microtubules in macrophage-like cells is reported to result in a rapid disassembly of podosomes ^34,35^.

Since the effects of microtubule disruption on focal adhesions and podosomes are contrary, one can suggest that microtubules regulate these integrin adhesions using different effectors. In our study reported here we surprisingly found a common mechanism underlying the effect of microtubules on both focal adhesions and podosomes. Firstly, we found that KANK family proteins are indeed involved in physical linking of microtubules to both focal adhesions and podosomes. Uncoupling of microtubules from integrin adhesions by interfering with KANK1 or KANK2 function or forced coupling by rapamycin-induced linking of KANK domains produced profound effect on both podosomes and focal adhesions. In particular, disconnection of microtubules from the integrin adhesions faithfully reproduced the contrary effects of microtubule disruption on these two structures. Next, we found that in both cases the effectors are myosin-II filaments, which are strongly augmented upon uncoupling microtubules from either podosomes or focal adhesions. Finally, such activation depends on the microtubule-associated RhoGEF, GEF-H1, which activates the Rho-ROCK signaling axis when microtubules are disconnected from integrin adhesions. Thus, KANK and GEF-H1-dependent local regulation of myosin-II filaments assembly appears to be a general mechanism underlying the effects of microtubules on both classes of integrin-mediated adhesions.

## Results

### Disruption of microtubules or uncoupling them from integrin-based adhesions results in augmentation of focal adhesions and suppression of podosomes

We addressed the effect of perturbations of the microtubule network on two types of integrin-mediated adhesions of cells to extracellular matrix, focal adhesions and podosomes. Focal adhesions are characteristic of fibroblast-like cells and we chose to use the human fibrosarcoma cell line, HT1080, which has a well-developed system of focal adhesions associated with actin filament bundles known as stress fibers. For podosomes, we choose the human monocytic THP1 cell line, which upon treatment with transforming growth factor-beta 1 (TGFβ1) undergo differentiation into macrophage-like cells forming numerous podosomes^36^. We used THP1 cells treated with TGFβ1 in all the experiments presented in this paper. In addition, we used cells of human colon adenocarcinoma line (HT-29) and human umbilical vein endothelial cells (HUVECs) as other examples of cells forming focal adhesions, and cells of murine macrophage RAW 264.7 line stimulated by phorbol 12-myristate 13-acetate (PMA) as podosome-forming cells.

To depolymerize microtubules, we used 1 μM nocodazole. Such treatment disrupted the microtubules completely in less than 15 minutes in both HT1080 and THP1 cells. To uncouple microtubules from adhesion structures in a more physiological manner, we employed recent findings that modular proteins of the KANK family (in particularly KANK1 and KANK2) physically connect microtubule tips with focal adhesions ^37^. Indeed, KANKs are known to interact with talin (a major integrin-associated protein localized to focal adhesions and podosome rings) and, via α and β liprins, with the membrane microtubule-targeting complex (ELKS, LL5β)^37^, which in turn bind to CLASP end-tracking proteins ^38^. In addition, KANK proteins can bind directly to the microtubule plus-end motor KIF21A ^37,39^ and, at least in Drosophila, to EB1 ^26^. We then compared the effects of total microtubule disruption with those caused by depletion of KANK1 or KANK2.

In agreement with previous studies, we found that disruption of microtubules led to an increase of focal adhesion number and size in HT1080 cells (Figure 1A and 2A), but completely disrupted podosomes in TGFβ1-stimulated THP1 cells (Figure 1B, 2B and C).

**Figure 1.**
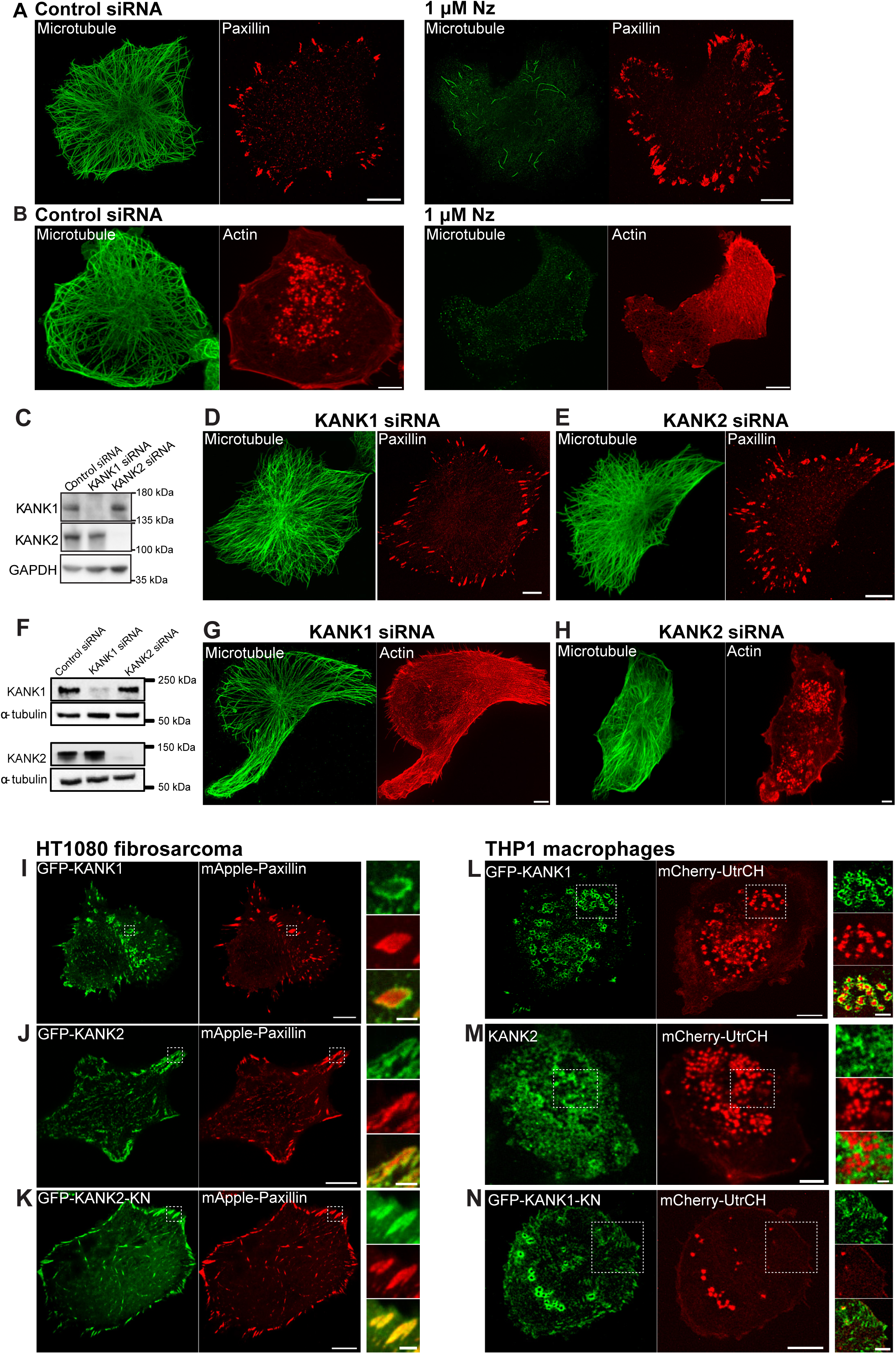
KANK family protein knockdowns or displacement mimic the effects of total microtubule disruption on focal adhesions and podosomes. (A) HT1080 cell transfected with control siRNA showed radially-organized microtubules (green) and numerous focal adhesions (red) at the cell periphery (left panel). Addition of 1 μM nocodazole disrupted the majority of microtubules and induced increase in focal adhesion size (right panel). (B) THP1 cells transfected with control siRNA and stimulated with TGFβ1 displayed radial microtubule organization (green) and numerous podosomes (red) (left panel). Addition of 1 μM nocodazole disrupted the microtubules and induced disassembly of podosomes in THP1 cells (right panel). (C) Western blots showing KANK1 and KANK2 levels in HT1080 cells. GAPDH was used as a loading control. (D, E) Knockdowns of KANK1 (D) or KANK2 (E) in HT1080 cells induced increase in focal adhesion size as compared to control (A, left panel) but did not interfere with microtubule integrity. (F) Western blots showing KANK1 and KANK2 levels in THP1 cells. α-tubulin was used as a loading control. (G, H) Knockdowns of KANK1 (G) but not KANK2 (H) in THP1 cells induce disassembly of podosomes without disruption of microtubules. Microtubules were visualized by immunostaining with α-tubulin antibody in both HT1080 and THP1 cells; focal adhesions in HT1080 cells were marked by immunostaining with paxillin antibody, while podosomes in THP1 cells were visualized by phalloidin to label F-actin cores. (I, J) Localization of GFP-KANK1 (I) and GFP-KANK2 (J) in HT1080 cells (green) as compared with focal adhesions labeling by expression of mApple-paxillin (red). Note that both KANK1 and KANK2 are often localized to the rim surrounding individual focal adhesions. For (I-N), the higher magnification images of the boxed areas and their merges are shown at the right. (K) HT1080 cell overexpressing the talin-binding domain of KANK2 (GFP-KANK2-KN, green) exhibited large focal adhesions (red). Note that GFP-KANK2-KN and mApple-paxillin have similar localizations. (L, M) Localization of GFP-KANK1 (L) and GFP-KANK2 (M) in THP1 cells (green) as compared with podosome labeling by mCherry-UtrCH (red). While GFP-KANK1 was found to surround the actin cores of podosomes, the association of KANK2 with podosomes was less prominent. (N) THP1 cell overexpressing talin-binding domain of KANK1 (GFP-KANK1-KN, green) demonstrated reduced podosome number and formation of focal adhesion-like structures. Scale bars in (A, D, E and I-K), 10 μm; (B, G, H and L-N), 5 μm; magnified images in (I-N), 2 μm.

**Figure 2.**
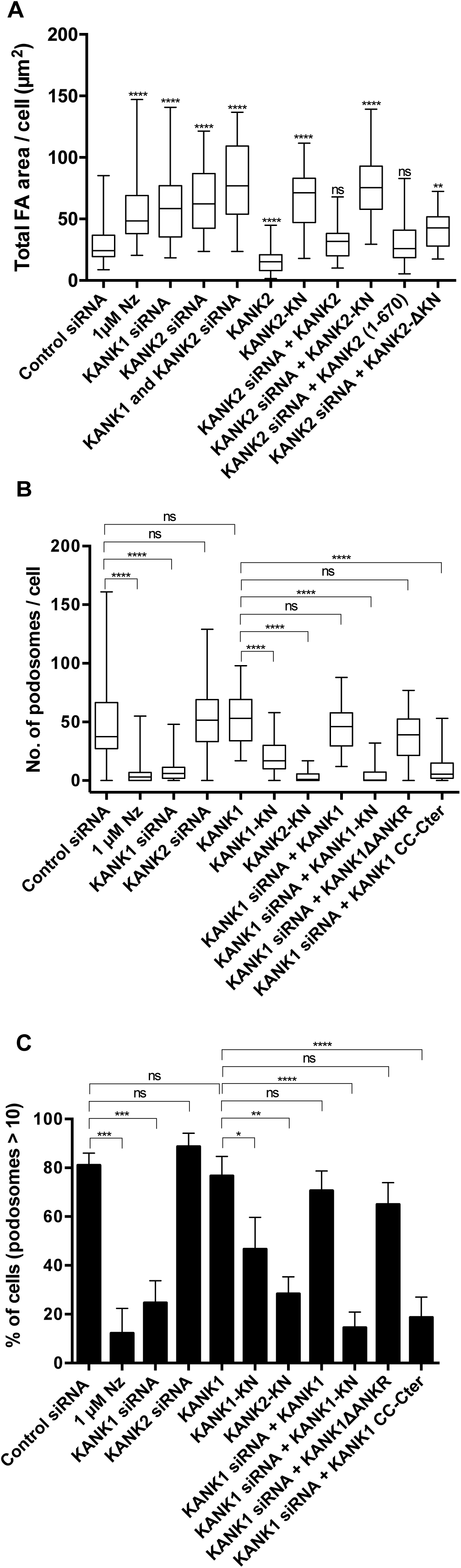
Evaluation of the effects of total disruption of microtubules and their disconnection from integrin adhesions on focal adhesion and podosomes. (A) The total areas of focal adhesions per cell (μm^2^) determined by measurement of HT1080 cells treated/transfected as indicated, after fixation and immunostaining with paxillin antibody. KANK2, KANK2-KN, KANK2 (1-670), ΚΑΝΚ2ΔΚΝ constructs (depicted in Figure 1I-K and Supplementary Figure 4H-M) were all tagged with GFP. (B, C) The numbers of podosomes per cell (B) and the percentage of cells with more than 10 podosomes (C) were determined in THP1 cells. Podosomes were visualized by mCherry-UtrCH in living cells transfected with GFP fusion constructs of KANK1, KANK1-KN, KANK2-KN, KANK1ΔANKR and KANK1-CC-Cter (depicted in Figure 1L, 1N, and Supplementary Figure 4B-G), or by phalloidin staining of fixed cells. The focal adhesion area and number of podosomes per cell are presented as box-and-whiskers plots, while the percentage of cells with more than 10 podosomes as mean ± SD. Nz-nocodazole. The significance of the difference between groups was estimated by two-tailed Student’s *t*-test, the range of p-values >0.05(non-significant), ≤ 0.05, ≤0.01, ≤0.001, ≤ 0.0001 are denoted by “ns”, one, two, three and four asterisks (*), respectively.

As shown by gene expression profiling (Supplementary Table 1), both KANK1 and KANK2 isoforms are expressed in the two cell lines used in this study, focal adhesion-forming HT1080 and podosome-forming THP1. Expression of two other isoforms, KANK3 and KANK4 was negligible. We confirmed previous observations on the localization of GFP-KANK1 and 2 around focal adhesion plaques ^28,37^ (Figure 1I and J). We further demonstrated that in THP1 cells stimulated by TGFβ1, GFP-KANK1 localize to the external ring surrounding podosomes (Figure 1L, Supplementary Figure 1B). Similar localization of KANK1 around the podosomes was also observed in PMA-stimulated RAW 264.7 macrophages (Supplementary Figure 2A). Association of KANK2 with podosomes in THP1 cell was less prominent than that of KANK1, even though a small number of podosomes were surrounded by KANK2-positive rings (Figure 1M).

In HT1080 cells, knockdown of both KANK1 and KANK2 (Figure 1C) produced an increase in focal adhesion size similar to total disruption of microtubules by nocodazole (Figure 1D, E and 2A). The double knockdown of KANK1 and KANK2 affected focal adhesions more strongly than the knockdown of KANK1 or KANK2 alone (Supplementary Figure 1C, Figure 2A). In THP1 cells, knockdown of KANK1 and KANK2 (Figure 1F) produced different effects. Only the knockdown of KANK1 induced pronounced disruption of podosomes in THP1 cells (Figure 1G, 2B and 2C), phenocopying the effect of total microtubule disruption. Knockdown of KANK2 (Figure 1H) did not significantly affect podosome number in THP1 cells (Figure 1H, 2B and 2C) even though it increased the cell projection area (Supplementary Figure 1A).

In addition to KANK1/2 knockdown, we uncoupled microtubules from integrin adhesions by overexpression of KANK1/2 truncated constructs, GFP-KANK1-KN and GFP-KANK2-KN, which bind to talin but not to microtubules ^28,37^. We anticipated that these constructs could compete with endogenous KANK1 and 2 for binding to talin. Indeed, expression of either GFP-KANK1-KN (Supplementary Figure 1D and E) or GFP-KANK2-KN (Figure 1K) in HT1080 cells resulted in localization of these constructs to focal adhesions and elimination of endogenous KANK2 from this location (Supplementary Figure 1D). Thus, we concluded that overexpression of GFP-KANK1/2-KN disconnects the focal adhesions from microtubules and examined how such treatment will affect focal adhesions. In HT1080 cells, overexpression of GFP-KANK2-KN led to an increase in focal adhesion size (Figure 1K and 2A) to a similar degree as KANK2 knockdown or nocodazole treatment. An increase in focal adhesion area upon overexpression of GFP-KANK2-KN was also observed in HT-29 epithelial cells. The effect in this case also mimicked the effect of nocodazole treatment (Supplementary Figure 3A-D). Finally, the overexpression of GFP-KANK1-KN significantly increased the focal adhesion area in endothelial cells (HUVECs) (Supplementary Figure 3E-G).

**Figure 3.**
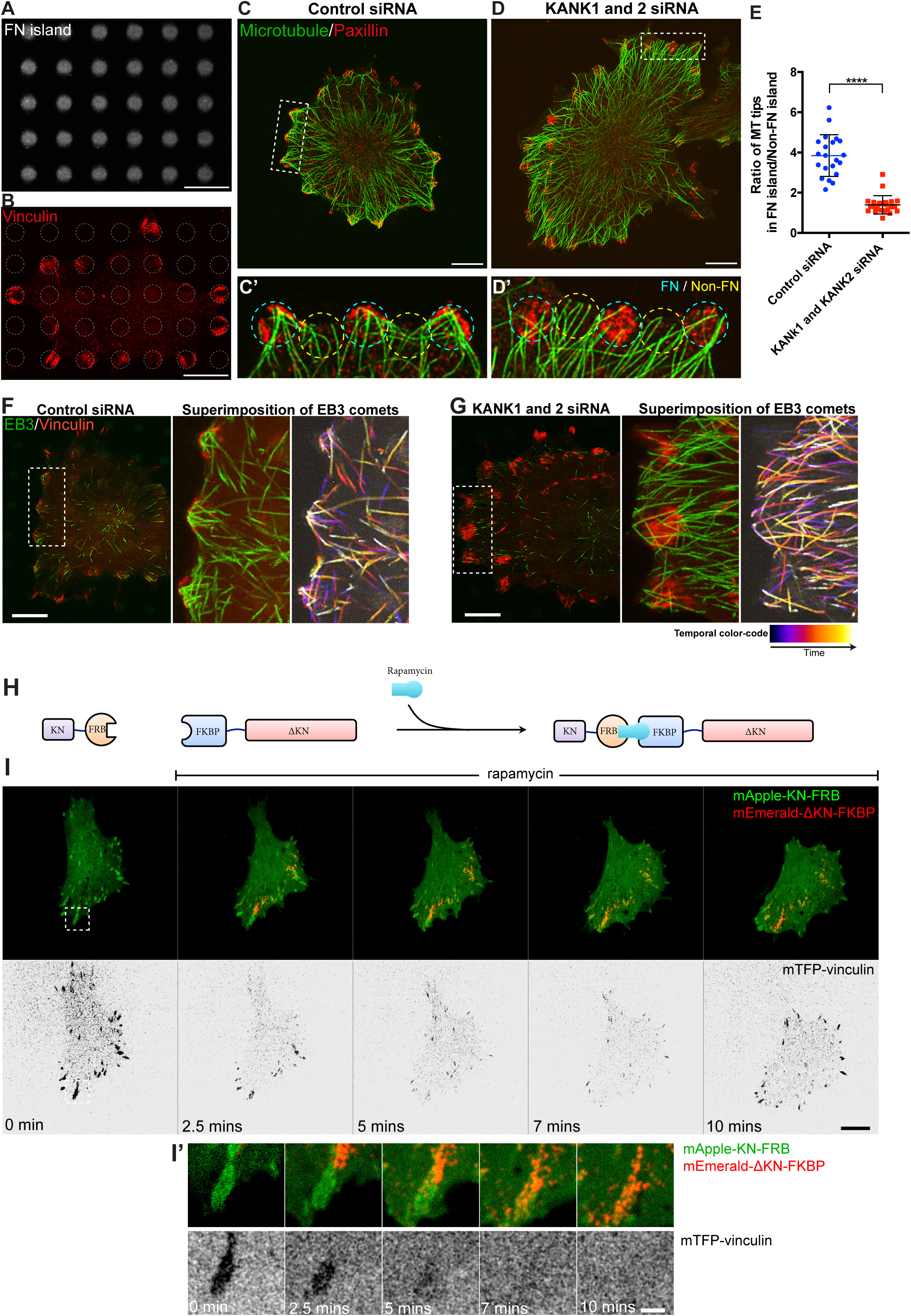
Depletion of KANK family proteins abolishes targeting of microtubules to focal adhesion in HT1080 cells (A) Micropatterned substrate used for ordered localization of focal adhesions. The adhesive fibronectin islands have diameter of 4 μm and arranged in square lattice with the period of 8 μm. (B) Immunostaining of focal adhesions by vinculin antibody (red) in HT1080 cell on micropatterned substrate (grey dotted circles). (C and D) Microtubules (green) and focal adhesions decorated by paxillin staining (red) in HT1080 cells on micropatterned substrate. In control cell (C), microtubule tips are concentrated at the areas occupied by focal adhesion, while in cells with double KANK1 and KANK2 knockdown, they are more randomly distributed. The boxed areas in C and D are shown at higher magnification in C’ and D’, respectively. The cyan dotted circles in C’ and D’ correspond to adhesive islands where focal adhesions are located. For the sake of comparison, the areas at the cells edge in between the adhesive islands were also marked (yellow dotted circles). Scale bars, 10 μm. (E) The ratios of the numbers of microtubule tips at z=0 overlapping with adhesive islands containing focal adhesions (as marked by cyan circles in C’ and D’) to the numbers of microtubule tips in non-adhesive regions of the same cell (as marked by yellow circles in C’ and D’). Each dot represents the value of such ratio for individual cell (7-13 islands per cell were assessed). Note that the difference between the number of microtubule tips overlapping with adhesive and non-adhesive islands is larger in control than in KANK1 and KANK2-depleted cells. (F and G) Microtubule targeting to focal adhesions in living control (F) and KANK1+KANK2 depleted (G) HT1080 cells. Microtubule tips were decorated by mKO-EB3 (green), focal adhesions were labeled by GFP-vinculin (shown in red). Left images in F and G panels show typical frames from time-lapse Movies S1 and S2, respectively. The central and right panels correspond to superimposition of 60 frames taken with 1-second interval. In the right images of F and G, the EB3 comets taken at different time intervals are marked according to temporal color-code (far right). Scale bars, 10 The significance of the difference between groups was estimated by two-tailed Student’s *t*-test, the range of p-values >0.05(non-significant), ≤ 0.05, ≤0.01, ≤0.001, ≤ 0.0001 are denoted by “ns”, one, two, three and four asterisks (*), respectively. (H, I and I’) Transient disassembly of focal adhesions upon the enforced targeting of microtubules. (H) A scheme depicting rapamycin-mediated formation of a link between the constructs corresponding to two parts of KANK1 molecule (mApple-KN-FRB and FKBP-ΔKN-mEmerald). (I) The cell triple transfected with mApple-KN-FRB (green), FKBP-ΔKN-mEmerald (red) and mTFP-vinculin (inverted black and white images) was incubated with rapamycin at 1 μg/ml. Note the localization of mApple-KN-FRB shown in green to focal adhesions before rapamycin addition. Rapamycin promoted the accumulation of FKBP-ΔKN-mEmerald to the focal adhesions which triggered their transient disassembly manifested by disappearance of vinculin. After 10 mins of incubation, focal adhesions re-assembled again. Scale bar, 10 μm. (I’) High magnification of the area boxed in (I). Note the details of localization of KN and ΔKN parts of KANK1 protein during the process of rapamycin linking and focal adhesion disassembly. Scale bar, 2 μm. See Movie S4.

Along the same line of reasoning, we overexpressed GFP-KANK1-KN in THP1 cells and found that such treatment led to a significant reduction in podosome number, resembling the effects of KANK1 knockdown or nocodazole treatment (Figure 1N, 2B, 2C). Exogenous GFP-KANK1-KN was localized to the few residual podosomes. Interestingly, the overexpression of talin-binding KANK2 domain (GFP-KANK2-KN) resulted in podosome disruption in THP1 cells (Supplementary Figure 1F, Figure 2B and 2C) in spite of the fact that knockdown if KANK2 did not disrupt podosomes. This suggests that the KN domains of KANK1 and KANK2 are similar and compete with each other for binding to talin.

Disappearance of podosomes in THP1 cells is sometimes accompanied by the appearance of focal adhesion-like structures. Such an effect is also prominent in another podosome-forming cell type, RAW 264.7 macrophages. Upon PMA stimulation, these cells form very prominent podosomes surrounded by GFP-KANK1-positive rings (Supplementary Figure 2A). Overexpression of GFP-KANK1-KN in these cells led to complete elimination of podosomes and formation of numerous elongated KANK1-KN-containing focal adhesion-like structures at the cell periphery (Supplementary Figure 2B-D).

To further investigate the mechanism of KANK1/2 effects, we performed rescue experiments on KANK1-depleted THP1 cells and KANK2-depleted HT1080 cells using various deletion mutants of KANK1 and KANK2, respectively (Supplementary Figure 4A). We found that constructs consisting of talin-binding domain alone (GFP-KANK1-KN, GFP-KANK2-KN), or constructs lacking a talin-binding domain (GFP-KANK1-CC-Cter, GFP-KANK2ΔKN), which cannot link talin-containing structures with microtubules, did not rescue the effects of KANK1 knockdown on podosomes and KANK2 knockdown on focal adhesions (Figure 2A-C, Supplementary Figure 4 D, E, J and K). At the same time, the constructs (GFP-KANK1ΔANKR, GFP-KANK2 (1-670)) containing the talin-binding domain together with the coil-coiled liprin-binding domain ^28,37,39^ rescued the effects of KANK1 and KANK2 knockdowns respectively, similarly to the full-length KANK1 or KANK2 (Figure 2A-C, Supplementary Figure 4B, C, H and I). Of note here, the c-terminal region of KANK1 missing from GFP-KANK1ΔANKR and GFP-KANK2 (1-670), which is known to be involved in binding KIF21A kinesin ^39^, appeared to be dispensable for the KANK effects on podosomes (Supplementary Figure 4F) and focal adhesions (Supplementary Figure 4L).

**Figure 4.**
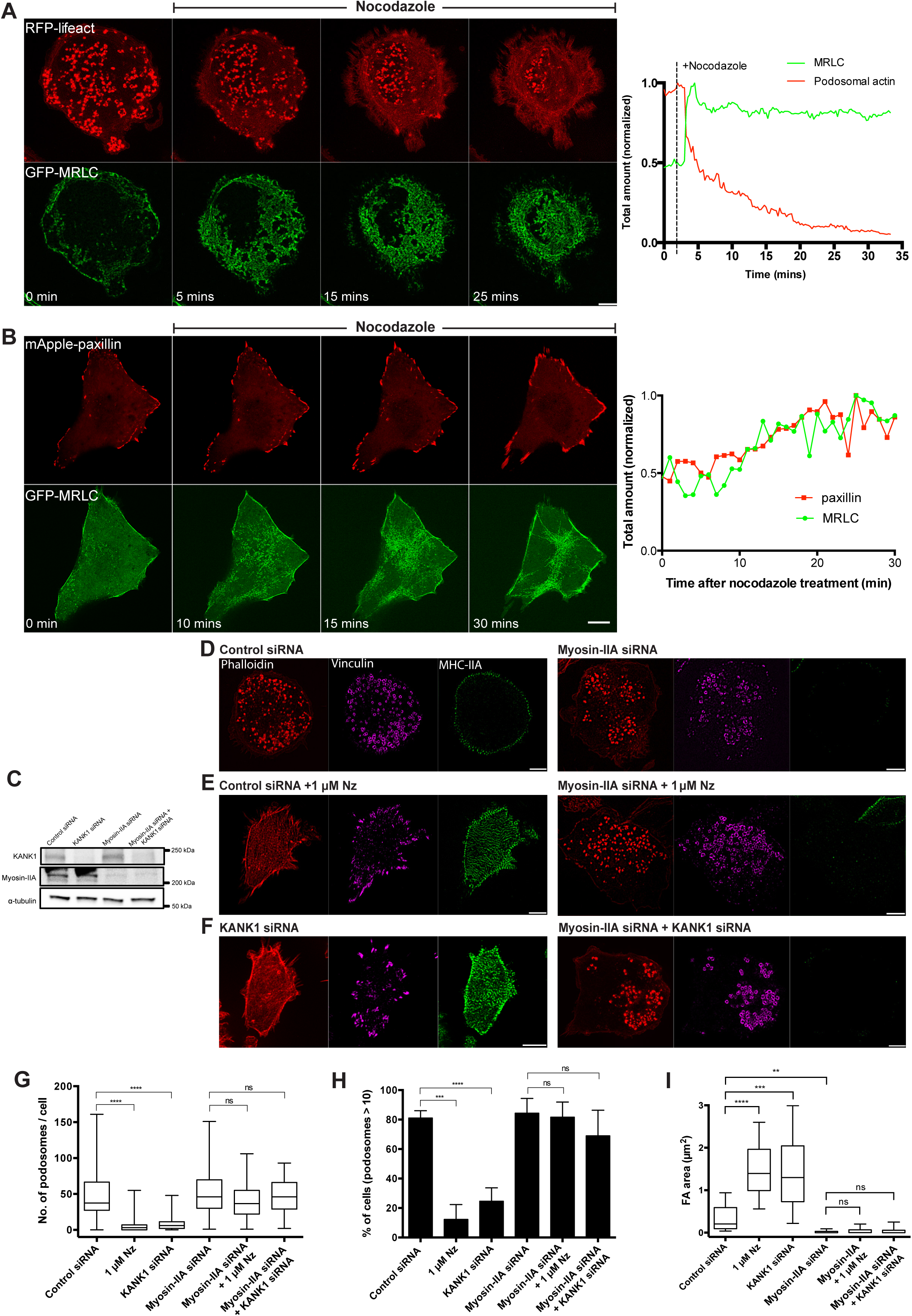
Myosin-II filaments mediate the effect of microtubules on integrin-based adhesions. (A and B) Addition of 1 μM nocodazole induced the assembly of numerous myosin-II filaments (green) in (A) THP1 and (B) HT1080 cells, which correlates with the disruption of podosomes (red) in THP1 cells (A) and increase in focal adhesion size (red) in HT1080 (B), respectively. Scale bars of each panel in (A) and (B) are 5 and 10 μΜ, respectively. Graphs on the right shows that upon microtubule disruption by nocodazole, the increase in the amount of myosin-II filaments was accompanied by both the decrease in podosome number in THP1 cell (A, upper row) and the increase in focal adhesion area in HT1080 cells (B, upper row). See Movies S5 and S6 for both THP1 and HT1080 cells, respectively. Focal adhesions were marked by expression of mApple-paxillin in HT1080 cells, while podosome actin cores and other F-actin structures were visualized by expression of RFP-lifeact in THP1 cells. Details of the quantification procedures can be found in the Materials and Method section. (C) Western blots showing KANK1 and myosin-IIA levels in TGFβ1-stimulated THP1 cells transfected with scramble (control), KANK1, or MYH9 siRNAs, as well as KANK1 and MYH9 siRNAs together. α-tubulin was used as a loading control. (D) Left panel: THP1 cells transfected with scrambled control siRNA showed circumferential assembly of myosin-II filaments (green) and numerous podosomes visualized by actin (red) and vinculin (purple) staining. Scale bars, 5μm. Boxed area shows higher magnification of small focal complexes and podosomes in the insets. Scale bars, μm. Right panel: Knockdown of MYH9 led to complete disappearance of myosin-IIA filaments but did not affect podosomes (actin: red; vinculin: purple). (E and F) Left panels: Microtubule disruption (E) or their uncoupling from podosomes by endogeneous KANK1 depletion (F) induced massive assembly and alignment of myosin-II filaments (green) as well as disruption of podosomes (red, purple) and formation of focal adhesion-like structures marked by vinculin (purple) staining. Scale bars, 5μm. Boxed area shows higher magnification of large focal adhesion-like structures in the insets. Scale bars, μm. Right panels: THP1 cells lacking myosin-IIA and treated with nocodazole (E) or with double knockdown of myosin-IIA and KANK1 (F) preserved the podosomes and do not form focal adhesions (actin:red; vinculin: purple). Scale bars, 5μm. (G-I) Quantification of the number of podosomes per cell (G), percentage of cells forming more than 10 podosomes (H) and area of focal adhesions (I) in THP1 cells treated as indicated in the previous figures (D-F). Nz-nocodazole. The significance of the difference between groups was estimated by two-tailed Student’s *t*-test, the range of P-values >0.05(non-significant), ≤ 0.05, ≤0.01, ≤0.001, ≤ 0.0001 are denoted by “ns”, one, two, three and four asterisks (*), respectively.

To better understand why KANK2 unlike KANK1 is poorly localized to podosomes, and dispensable for their integrity, we examined the role of the liprin-binding coiled coil domain of KANK1 in podosome regulation. We created a chimeric construct of KANK2 (KANK2/CC-KANK1) in which its own coiled-coil domain was substituted with that of KANK1 (Supplementary Figure 5A). Such chimeric construct localized to podosome rings much better than KANK2. Moreover, while wild-type KANK2 did not rescue podosomes in KANK1-depleted cells, the GFP fusion of KANK2/CC-KANK1 significantly rescued podosomes in the absence of KANK1 (Supplementary Figure 5B-F). This suggests that KANK2 has a lower affinity to podosomes in THP1 compared to KANK1 because of deficient liprin-binding and therefore cannot appropriately connect podosomes with microtubules in these cells.

**Figure 5.**
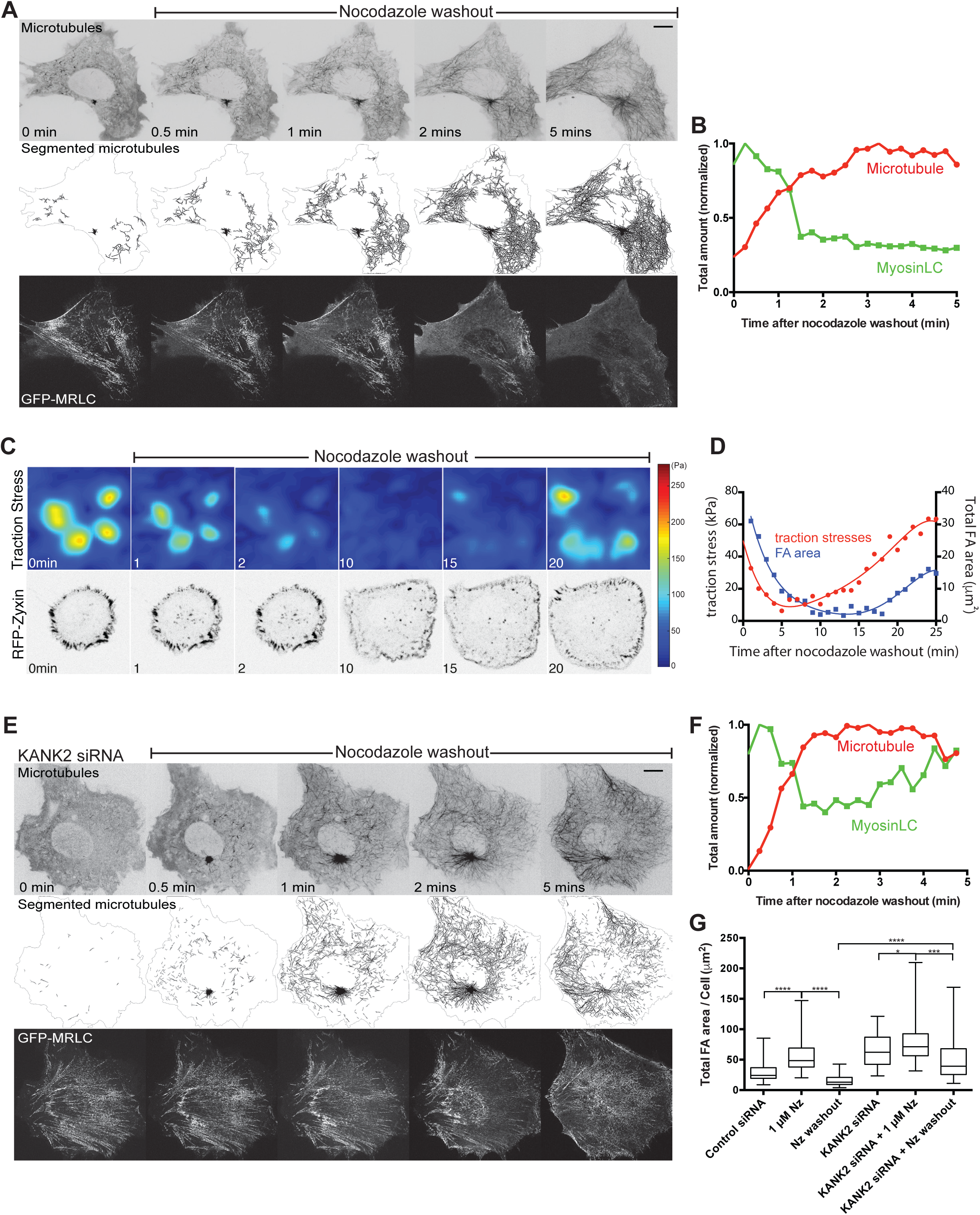
Augmentation of microtubule contacts with focal adhesions brings about disassembly of myosin-II filaments and focal adhesions. (A) Time-lapse observations of microtubules (upper row), segmented microtubules (second row) using open-source software package called SIFNE ^92^ and myosin-II filaments (bottom row) in the course of microtubule outgrowth after nocodazole washout in HT1080 cell. Time after the nocodazole washout is indicated in each frame. Microtubules and myosin-II filaments were labeled by mApple-MAP4 and GFP-MRLC, respectively. Note that recovery of microtubules is accompanied by disappearance of myosin-II filaments. See Movie S7. (B) Quantification of the changes in the amount of microtubules (red) and myosin-II filaments (green) shown in (A). Details of the quantification procedures can be found in the Materials and Method section. (C) Changes in traction forces exerted by HT1080 cells in the course of microtubule recovery following nocodazole washout. The traction forces are presented in the upper row in spectrum scale (right). The corresponding images of cells with focal adhesions labeled with RFP-zyxin are presented in the bottom row, along with the time in minutes after nocodazole washout. Note that the traction forces and focal adhesion size began to decrease already in 1 minute following nocodazole washout but recover in about 20 minutes. See Movie S8. (D) Changes in sum of the absolute values of traction forces (kPa) and total focal adhesion area (μm^2^) per cell shown in (C). Note that drop and subsequent increase of traction forces (red) preceded the drop and increase in focal adhesion area (blue), respectively. (E) KANK2 knockdown in HT1080 suppresses the effect of microtubule outgrowth on the amount of myosin-II filaments. Microtubules (upper row), segmented microtubules (second row), myosin-II filaments (bottom row) in the same cell were labeled as indicated in (A). Time after nocodazole washout is indicated in the upper row. See Movie S9. Note that in spite of microtubule outgrowth, the changes in myosin-II filaments were less pronounced than in control cell shown in (A). (F) Quantification of the changes in amount of microtubules (red) and myosin-II filaments (green) was performed as in (B). (G) Changes in the total area of focal adhesions after nocodazole treatment and washout in control and KANK2 knockdown HT1080 cells. The cells before and after corresponding treatments were fixed and stained with paxillin antibody to visualize focal adhesions. Nz-nocodazole. Not less than 40 cells were measured for each type of treatment. The data presented as box-and-whiskers plots and statistical significance of the difference (p-values) was calculated by two-tailed Student’s *t*-test, the range of P-values >0.05(non-significant), ≤ 0.05, ≤0.01, ≤0.001, ≤ 0.0001 are denoted by “ns”, one, two, three and four asterisks (*), respectively. Scale bars in all panels correspond to 10 μm.

### KANK proteins control focal adhesion by physically connecting them with microtubules

In view of strong effect of KANK family protein depletion on focal adhesions that mimics complete disruption of microtubules, we decided to directly assess the effect of KANK protein knockdown on microtubule targeting to focal adhesions in HT1080 cells. In order to better visualize the microtubule targeting to focal adhesions, we plated cells on to patterned substrata consisting of small adhesive islands (diameter 4 μm) organized into square lattice with a periodicity of 8 μm (Figure 3A). The cells spreading on such substrates form focal adhesions only within the adhesive islands (Figure 3B). In control cells, microtubule tips are concentrated at such islands which can be seen by examining fixed specimens after immunostaining of microtubules and focal adhesions (Figure 3C, C’ and E), as well by live imaging of microtubule tips by expression of mKO-EB3-end tracking protein along with GFP-vinculin as a focal adhesion marker (Figure 3F, Movie S1). In the cells depleted of KANK1 and KANK2, the focal adhesions were larger than in control cells (c.f. Figure 2A and Supplementary Figure 1C), and the microtubule tips were more randomly distributed, filling almost evenly the space between the adhesive islands (Figure 3D, D’ and G, Movie S2), with only some slight preference to focal adhesions (Figure 3E). We will demonstrate the effect of KANK1 depletion on the targeting of microtubules to podosomes in Figure 7 below.

**Figure 6.**
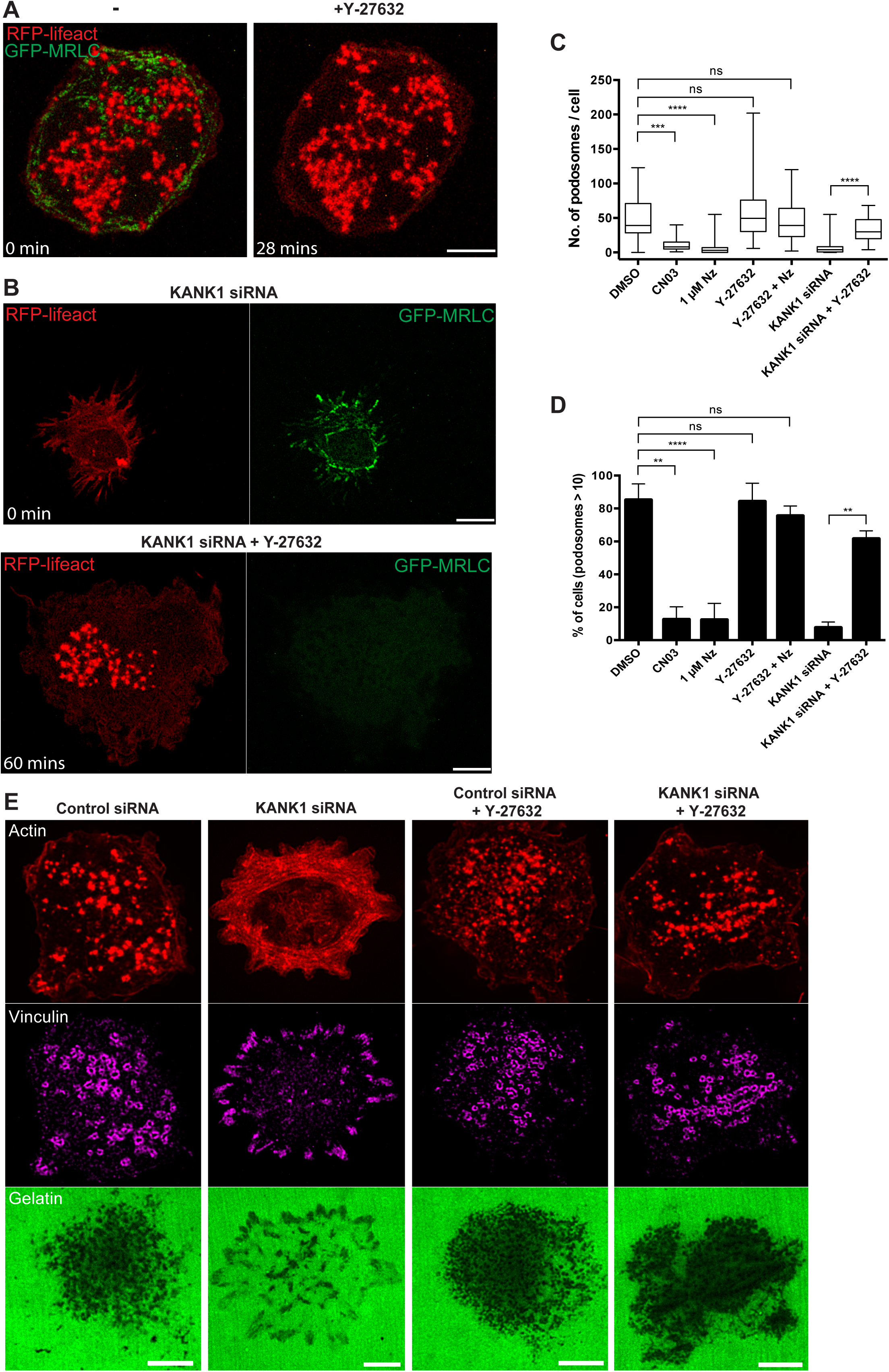
Inhibition of ROCK results in recovery of functional podosomes in KANK1-depleted cells. (A) Podosomes (red) and myosin-II filaments (green) in THP1 cell before (left) and after (right) addition of ROCK inhibitor, 30 μΜ Y-27632. Note that Y-27632 treatment totally disrupted myosin-II filaments (green) but not podosomes (red). Scale bar, 5 μΜ. See Movie S12. (B) KANK1-depleted THP1 cell shown before (upper panel, t=0) and after (lower panel, t=60 mins) incubation with 30 μΜ Y-27632. Note the disappearance of myosin-II filaments (green) and recovery of podosomes (red) upon Y-27632 treatment. Scale bars, 5 μm. See Movie S15. (C and D) The number of podosomes (C) and the percentage of cells with more than 10 podosomes (D) in THP1 cells treated as indicated. CN03 is a Rho activator, Nz - nocodazole, Y-27632 is a ROCK inhibitor. See images illustrating all treatments in Figure 6A, 6B and Supplementary Figure 9A-D. “Y-27632+Nz” represents the pooled results of experiments shown in Supplementary Figure 9C and 9D. The data presented as box- and-whiskers plots and statistical significance of the difference (p-values) was calculated by two-tailed Student’s *t*-test, the range of P-values >0.05(non-significant), ≤ 0.05, ≤0.01, ≤0.001, ≤ 0.0001 are denoted by “ns”, one, two, three and four asterisks (*), respectively. (E) Podosomes are visualized by actin (red, upper row) and vinculin (purple, middle row) staining, while matrix degradation by the same cells (lower row) is seen as dark areas in the fluorescent gelatin substrates (green). KANK1 depletion resulted in disappearance of podosomes and formation of focal adhesions (second column from the left), which have reduced ability to degrade fluorescent gelatin substrate. Treatment with Y-27632 does not affect substrate degradation by podosomes in control cell (third column) and recovered functional podosomes in KANK1 knockdown cell (fourth column). Scale bars, 5 μm.

**Figure 7.**
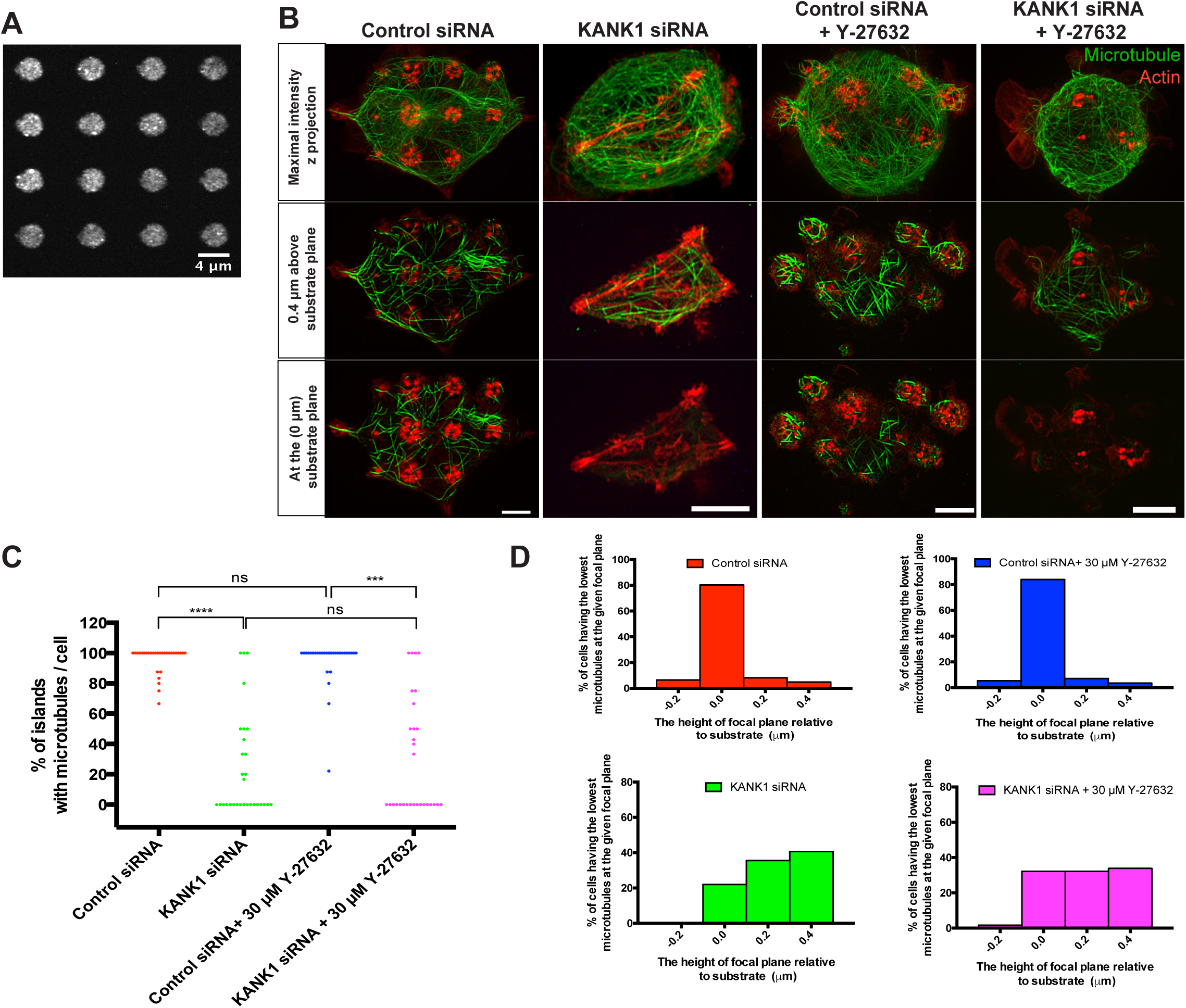
Depletion of KANK1 abolishes targeting of microtubules to podosomes in THP1 cells. (A) Micropatterned substrate used for ordered localization of focal adhesions. The adhesive fibronectin islands have diameter of 4 μm and arranged in square lattice with the period of 8 μm. (B) Microtubules visualized by α-tubulin antibody staining (green) and actin labeled by phalloidin (red) in THP1 cells plated on micropatterned substrate. Upper row shows z projection of all microtubules and actin structures; middle and lower rows show the microtubules and actin in the same cell at the focal planes located 0.4 μm above the substrate and at the substrate plane (z= 0.0 μm). In control cell (first column from the left), the actin decorates podosomes located at the adhesive islands. KANK1 knockdown (second column) resulted in the disruption of podosomes and appearance of focal adhesion-like structures. Y-27632 treatment does not affect podosomes in control cell (third column) and rescued them in KANK1 knockdown cell (fourth column). Note that microtubules overlapped with podosomes at the z=0 focal plane, in cells transfected with control siRNA but not in cells transfected with KANK1 siRNA. Scale bars, 5μm (C and D) Quantification of the overlap of microtubule tips with the adhesive islands containing podosomes or focal adhesion-like structures. (C) Percentage of such adhesive islands containing the microtubule tips at z=0 focal plane. Each dot corresponds to individual cell (n=30 cells were scored for each type of treatment). The statistical significance of the difference (p-values) was calculated by non-parametric Mann-Whitney U test, the range of P-values >0.05(non-significant), ≤ 0.05, ≤0.01, ≤0.001, ≤ 0.0001 are denoted by “ns”, one, two, three and four asterisks (*), respectively. (D) The histograms representing fractions of the cells having the lowest microtubule tips at indicated focal planes (n≥56 cells were scored for each type of treatment).

While disruption of microtubules resulted in augmentation of focal adhesions, microtubule outgrowth after nocodazole washout led to transient disassembly of focal adhesions ^20^. We reproduced the effect of microtubule outgrowth on focal adhesions in HT1080 cells (Movie S3). In order to elucidate the role of KANK proteins in this process, we used the technique of rapamycin-induced dimerization to link two parts of KANK1 protein: the KN talin-binding domain and the rest part of the molecule (ΔKN) (Figure 3H). Transfection of HT1080 cells with mApple-KN-FRB and FKBP-ΔKN-mEmerald resulted in an increase of focal adhesion size since mApple-KN-FRB, similarly to other KN constructs described above (Supplementary Figure 1D), apparently displace endogenous KANKs from talin. The addition of rapamycin which links mApple-KN-FRB with FKBP-ΔKN-mEmerald led to accumulation of FKBP-ΔKN-mEmerald to KN-containing adhesions and subsequently triggered transient disassembly of focal adhesion manifested by disappearance of vinculin (Figure 3I and I’, Movie S4). Thus, enforced targeting of KANK1-ΔKN involved in microtubule capturing to the KANK1-KN localized to focal adhesions mimicking the effect of microtubule outgrowth.

### Myosin-II filaments mediate the effect of microtubules on integrin-based adhesions

We next visualized myosin-II filaments by imaging either GFP-tagged myosin-II regulatory light chain (GFP-MRLC) or myosin-IIA heavy chain antibody by super-resolution structured illumination microscopy (SIM). We observed that disruption of microtubules resulted in massive formation of new myosin-II filaments. This effect was especially dramatic in THP1 cells, which normally contain only a rim of myosin-II filaments associated with circumferential actin bundles at the cell periphery (Figure 4A, Movie S5). Upon addition of nocodazole, a rapid disassembly of podosomes occurs in parallel with a drastic increase in the number of new myosin-II filaments (Figure 4A, graph on the right, Movie S5). Approximately 1 hour following nocodazole addition, the newly formed myosin-II filaments aligned in registry forming well-developed myosin-II filament stacks (Figure 4A, 4E and Movie S6). This process was accompanied by formation of short actin filament bundles, some of which were associated with newly formed focal adhesion-like structures (Figure 4I). Even though HT1080 cells contained some myosin-II filaments associated with stress fibers prior to nocodazole treatment (Figure 4B, Movie S6), the disruption of microtubules significantly increased the amount of myosin-II filaments as shown in Figure 4B. This increase, as in the case of THP1 cells, was accompanied by growth of focal adhesions. The epithelial HT-29 cells also responded to microtubule disruption by augmentation of myosin-II filaments (Supplementary Figure 3B) in agreement with ref ^40^.

To ascertain whether myosin-IIA filament assembly and disassembly are consequential or causal factors that account for the observed changes in focal adhesions and podosomes, we interfered with myosin-II filament formation by siRNA-mediated knockdown of myosin-IIA. Consistent with previous studies ^41,42^, we demonstrated that myosin IIA-depleted HT1080 cells essentially lack mature focal adhesions but contain small nascent adhesions at the cell periphery (Supplementary Figure 6A and B). In THP1 cells lacking myosin-IIA, podosome appearance did not differ from that seen in control cells (Figure 4D, G and H). However, in contrast to normal, these podosomes were insensitive to disruption of microtubules by nocodazole (Figure 4E, G and H). Thus, we conclude that depletion of myosin-IIA filaments abolished the effect of microtubule disruption on both focal adhesions and podosomes.

We observed that knockdown of KANK1 in THP1 cells also produced a significant increase of the number of myosin-IIA filaments (Figure 4F, left panel). In contrast to control cells (Figure 4D, left panel), in which myosin-II filaments were located only at the periphery, in KANK1-depleted THP1 cells, myosin-IIA filaments formed organized structures associated with bundles of actin filaments (Figure 4F, left panel). Many cells, similarly to nocodazole-treated cells, demonstrated formation of focal adhesions associated with these bundles (Figure 4F left panel and I). In HT1080 cells, KANK2 knockdown resulted in significant increase in formation of myosin-II filaments and focal adhesion area (Supplementary Figure 6C, middle panel).

Next, we assessed whether myosin-IIA depletion would interfere with the effects of KANK knockdown on podosomes and focal adhesions. We found that THP1 cells with double knockdowns of KANK1 and myosin-IIA demonstrated numerous podosomes, unlike the case of cells with knockdown of KANK1 alone (Figure 4F-H). Consistently, myosin-IIA knockdown in HT1080 cells permitted the formation of only nascent adhesions, abolishing the difference in focal adhesion size between KANK2-positive and KANK2 knockdown cells (Supplementary Figure 6C, middle and right panels). Thus, KANK-dependent modulation of both focal adhesions and podosomes is mediated via regulation of myosin-IIA filament assembly. Unlike myosin-IIA, the knockdown of myosin-IIB did not prevent the effects of microtubule disruption on both focal adhesions and podosomes. Interestingly, the knockdown of myosin-IIB in HT1080 cells by itself induced some increase in formation of myosin-IIA filaments and the growth of focal adhesions (Supplementary Figure 6D-F). However, in the myosin-IIB-depleted HT1080 cells, disruption of microtubules still significantly enhanced the myosin-IIA filament level and induced further increase in focal adhesion size (Supplementary Figure 6E and F). Similarly, knockdown of myosin-IIB did not prevent augmentation of myosin-IIA-containing filaments and elimination of podosomes upon KANK1 knockdown in THP1 cells (Supplementary Figure 7A-E). The number and degree of organization of myosin-II filaments in cells with double knockdown of KANK1 and myosin-IIB were even higher than in cells with KANK1 knockdown alone (Supplementary Figure 7D).

Outgrowth of microtubules following nocodazole washout in HT1080 cells led to a decrease in myosin-II filaments in the first minutes (Figure 5A and graph 5B, Movie S7). We also performed traction force microscopy to characterize the myosin-II driven contractility of HT1080 cells upon microtubule outgrowth. The traction forces exerted by cells on a substrate decreased upon microtubule recovery (Figure 5C, D, and Movie S8), consistent with the reduction of myosin-II filaments number (Figure 5B) but 20 minutes following the washout returned to the control level. The drop and increase in traction forces preceded the decrease and increase in focal adhesion area, respectively (Figure 5C).

Finally, we checked how microtubule outgrowth affects myosin-II filaments and focal adhesions in KANK2 knockdown cells. We found that KANK2-depleted HT1080 cells still preserved higher levels of myosin-II filaments than control cells under conditions of microtubule outgrowth (Figure 5E, bottom row, and F, Movie S9). Accordingly, microtubule outgrowth in KANK2 knockdown cells reduced focal adhesion size in lesser degree microtubule outgrowth in control cells (Figure 5G). These data suggest that microtubule outgrowth affects focal adhesions and myosin-II filaments only if microtubules can interact with focal adhesions locally via KANK.

### KANK-dependent targeting of microtubules regulates podosomes and focal adhesions via modulation of the Rho-ROCK signaling axis

Since the major pathway regulating myosin-II filament assembly is phosphorylation of the myosin-II regulatory light chain ^14^, we examined how changes of activity of the upstream regulators of such phosphorylation, namely Rho kinase (ROCK) and its activator RhoA, affected the processes of microtubule-driven regulation of focal adhesions and podosomes. Microtubule disruption resulted in an increase in the fraction of RhoA-GTP in both HT1080 and THP1 cells (Supplementary Figure 8A and B) in agreement with previous studies ^43^. KANK1 and KANK2 knockdown in HT1080 cells also led to increased RhoA-GTP levels (Supplementary Figure 8A and A’). Consistent with the fact that only knockdown of KANK1 but not of KANK2 led to disruption of podosomes in THP1 cells, we found that KANK1 knockdown but not KANK2 knockdown led to an increase in RhoA-GTP levels in THP1 cells (Supplementary Figure 8B and B’).

**Figure 8.**
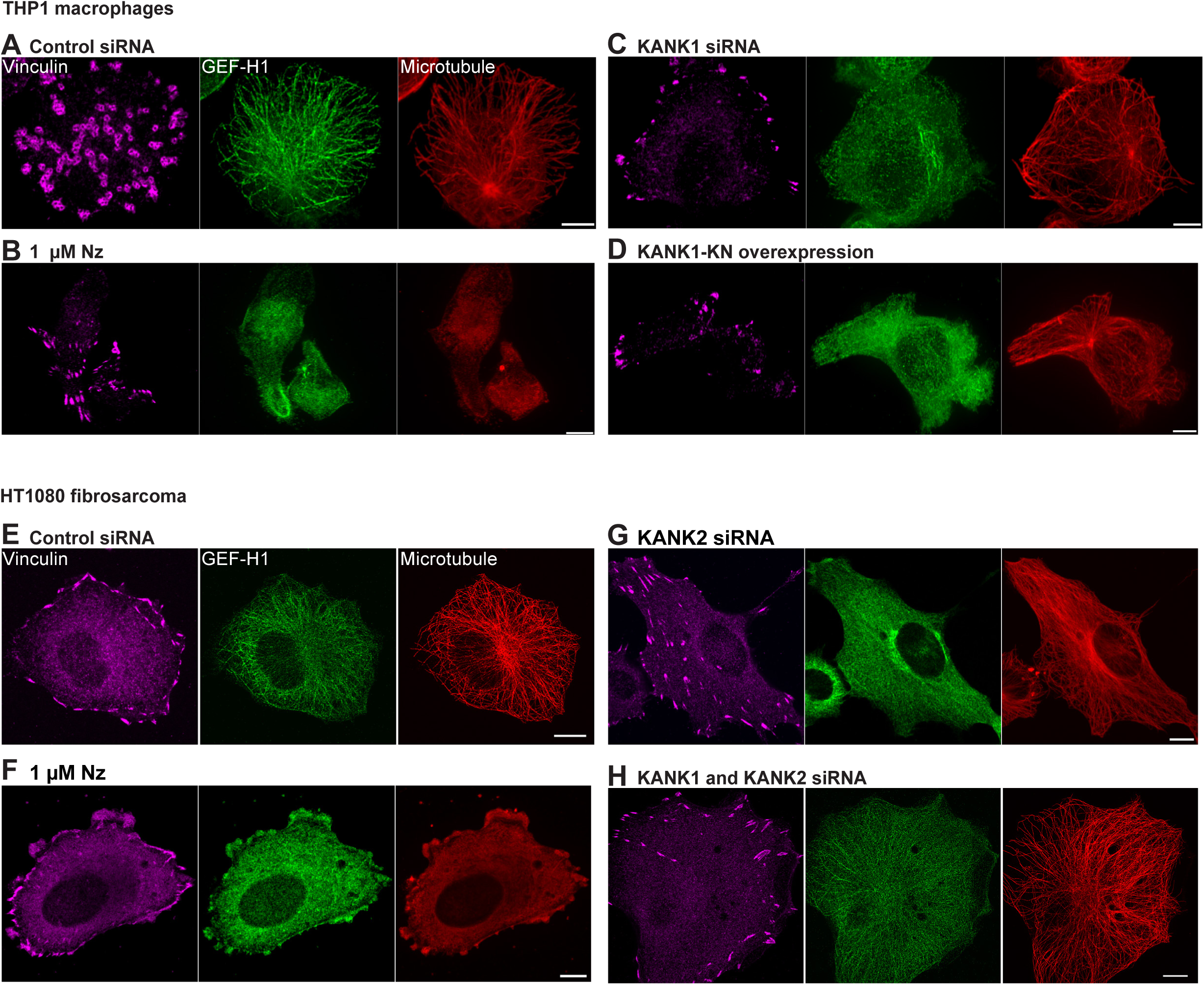
Uncoupling of microtubules from integrin adhesions results in release of GEF-H1 from microtubules in THP1 and HT1080 cells. Vinculin (purple), GEF-H1 (green) and α-tubulin (red) were visualized by immunofluorescence staining. (A) In control THP1 cell, vinculin labels podosomes, while GEF-H1 localize on microtubules. (B) Disruption of microtubules in THP1 cell by nocodazole treatment led to disappearance of podosomes and formation of focal adhesions marked by vinculin. The GEF-H1 is diffusely distributed over the cell. (C and D) Uncoupling of microtubules from talin-containing integrin adhesions in THP1 cells by KANK1 knockdown (C) or overexpression of talin-binding KN domain of KANK1 (D) led to disappearance of podosomes and formation of focal adhesions, similarly to total microtubule disruption. These treatments preserved microtubule integrity but induce release of GEF-H1 from microtubules. Scale bars, 5 μm. (E) In control HT1080 cell form peripheral focal adhesions labeled by vinculin; GEF-H1 is essentially localized to microtubules. (F) Disruption of microtubules in HT1080 cells resulted in augmentation of focal adhesions accompanied by release of GEF-H1 from microtubules. (G and H) KANK2 knockdown (G) or KANK1 and KANK double knockdown (H) in HT1080 cells resulted in augmentation of focal adhesions, preserves microtubule integrity but induces release of GEF-H1 from microtubules. Scale bars, 10 μ

We further investigated how pharmacological inhibition or activation of the Rho/ROCK pathway affected the processes of podosome disruption upon microtubule depolymerization and focal adhesion disruption upon microtubule outgrowth. We found that Rho activation by the small peptide CN03, mimicked the disruptive effect of microtubule disassembly or KANK1 depletion on podosomes (Figure 6C and D, Supplementary Figure 9A, Movie S10). Addition of DMSO (Supplementary Figure 9B, Movie S11) or Y-27632 (Figure 6A, Movies S12) did not affect podosome formation in control THP1 cells (Figure 6C and D). At the same time, inhibition of ROCK by Y-27632 was sufficient to prevent the podosome disruption normally seen after nocodazole treatment (Figure 6C and D, Supplementary Figure 9C, Movie S13). Moreover, addition of Y-27632 to THP1 cells incubated with nocodazole for several hours and completely lacking podosomes, still efficiently rescued podosomes by disassembling myosin-II filaments within 5-10 minutes (Figure 6C and D, Supplementary Figure 9D, Movie S14). Importantly, not only Y-27632, but also another specific ROCK inhibitor, thiazovivin ^44^, of different chemical nature, efficiently rescued the podosomes in the nocodazole-treated cell (Supplementary 10A-E).

**Figure 9.**
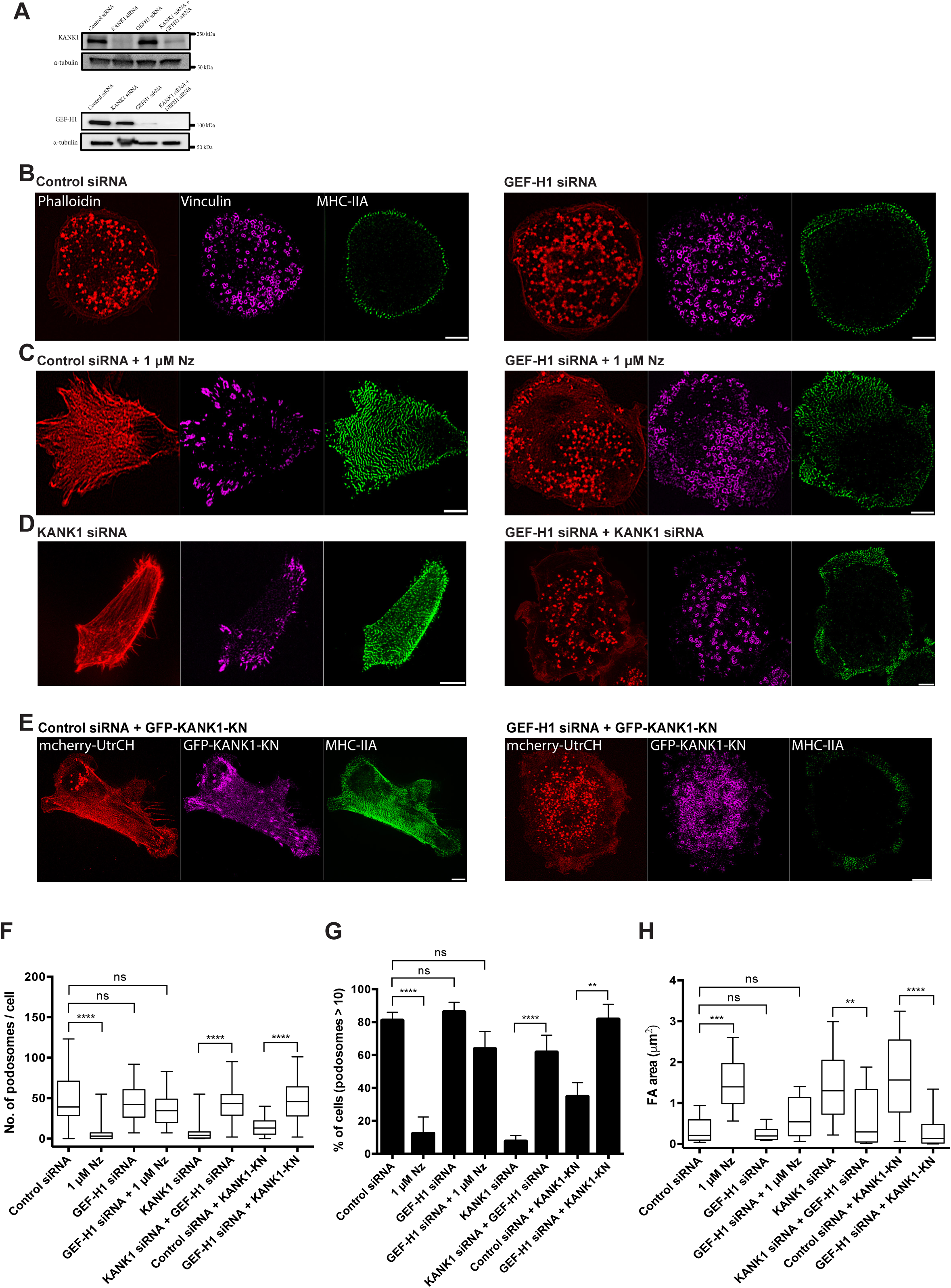
GEF-H1 is required for the microtubule-driven regulation of the myosin-II filaments, focal adhesions and podosomes. (A) Western blot demonstrating siRNA-mediated depletion of GEF-H1, KANK1, and both of them in THP1 cells. α-tubulin is used as loading control. (B) Visualization of podosomes (actin: red, vinculin: purple) and myosin-II filaments (green) in cells from cultures transfected with control siRNA (left panel) and GEF-H1 siRNA (right panel). The distribution of both podosomes and myosin-II filaments were unchanged in GEF-H1 knockdown cell. (C) Effect of microtubule disruption by on myosin-II filaments and podosomes in control (left panel) and GEF-H1 knockdown (right panel) cells. Labeling of myosin-II filaments and podosomes is the same as in (B). Note that increase in myosin-II filaments amount accompanied by disappearance of podosomes and appearance of vinculin-positive focal adhesions upon nocodazole treatment in control but not in GEF-H1 knockdown cells. (D) Podosomes, focal adhesions and myosin-II filaments in cells from KANK1 knockdown culture (left panel) and in cells from KANK1 and GEF-H1 knockdown culture. Labeling is the same as in (B) and (C). While KANK1 knockdown cell (left panel) demonstrate numerous myosin-II filaments and focal adhesions but not podosomes, the cells with KANK1/GEF-H1 double knockdown (right panel) preserved podosomes and did not show abundant myosin-II filaments. In (B-D), podosomes were visualized by phalloidin labeling of actin (red), and immunofluorescence staining of vinculin (purple). The myosin-II filaments in the same cells were labeled with MHCIIA antibody (green). (E) Disruption of podosomes by overexpression of GFP-KANK1-KN (left panel) is abolished in cells with knockdown of GEF-H1 (right panel). Actin was visualized by mCherry-UtrCH (red), GFP-KANK1-KN is shown in purple, and myosin-II filaments are shown in green. Scale bars for (B-E), 5 (F-H) Quantification of the number of podosomes (F), percentage of cells forming more than 10 podosomes (G) and average area of focal adhesions (H) in control THP1 cells and cells treated as shown in the previous figures (B-D). Nz=nocodazole. Not less than 80 cells were assessed for each type of treatment. The statistical significance of the difference (p-values) was estimated by two-tailed Student’s *t*-test.

Inhibition of ROCK by Y-27632 also led to rescue of podosomes in KANK1 knockdown THP1 cells (Figure 6B-D, Movie S15). Rescued podosomes were functionally active in terms of their ability to degrade extracellular matrix. The THP1 cell transfected with control siRNA demonstrated characteristic pattern of degradation of fluorescent gelatin (Figure 6E) in agreement with previous studies ^45^. The depletion of KANK1 induced elimination of podosomes and formation of focal adhesions, which were still capable of degrading the fluorescent gelatin (c.f. ref ^46^) but much more weakly than podosomes. Treatment of KANK1-depleted cells with Y-27632 not only rescued morphologically normal podosomes but also restore the gelatin degradation pattern (Figure 6E).

Experiments with podosome rescue in KANK1 knockdown cells by treatment with Y-27632 make it possible to check that KANK1 indeed is responsible for targeting of microtubules to podosomes. Similar to experiments with targeting of microtubules to focal adhesions described above (Figure 3), we plated the podosome-forming THP1 cells on micro-patterned substrate so that podosomes were formed only inside the small adhesive islands (Figure 7A and B). Confocal SIM of THP1 cells transfected with control siRNA, revealed an overlap of microtubule tips and podosomes at adhesive islands at the lowest focal plane (z= 0 μm). At the same time, in cells lacking KANK1, podosomes were absent and microtubule tips were not found at the z=0 μm plane, where focal adhesion-like structure were located (Figure 7B-D). Treatment with Y-27632 by itself affected neither podosome formation nor localization of microtubules to podosome-containing islands (Figure 7B). However, such treatment of KANK1 knockdown THP1 cells led to recovery of functional podosomes as already mention previously (Figure 6B-D and 7B). Important, the recovered podosomes unlike the control ones were not associated with microtubule tips (Figure 7B). The microtubule tips were not found at z=0 μm plane, where recovered podosomes were located. Quantification of the association of microtubule tips with podosomes at micro-patterned adhesive islands is given in Figure 7C-D. These data showed that KANK1 indeed is necessary for targeting of microtubules to podosomes. In addition, together with data in Figure 4 and Figure 6, these results indicate that functional podosomes can be maintained without a connection to microtubules if the Rho/ROCK-dependent myosin-II filament formation is suppressed.

Whilst disruption of podosomes is induced by disruption of microtubules, disruption of focal adhesion occurs upon microtubule outgrowth. We showed above that this process was preceded by the decrease of traction forces (Figure 5C) due to disassembly of myosin-II filaments (Figure 5A). Since myosin-II filament formation can be induced by activation of RhoA (Supplementary Figure 9A), we checked whether activation of the Rho/ROCK signaling axis would modulate the effect of microtubule outgrowth on the focal adhesions. Indeed, expression of either constitutively active RhoA (Q63L), or constitutively active ROCK mutant (rat Rok-alpha 1-543aa) in HT1080 cells prevented the disruptive effect of microtubule outgrowth on focal adhesions (Supplementary Figure 9E-G, Movie S16). Thus we conclude that microtubules regulate focal adhesions and podosomes through the same pathway modulating the level of Rho/ROCK activity.

### RhoGEF GEF-H1 is a major mediator of the microtubule effect on myosin-II filaments, focal adhesions and podosomes

The Rho nucleotide exchange factor, GEF-H1 is associated with microtubules in both THP1 and HT1080 cells (Figure 8) as well as HUVECs (Supplementary Figure 3I) in agreement with previous observations ^47,48,49^. Disruption of microtubules resulted in a diffuse distribution of GEF-H1 in all cell types (Figure 8B and 8F). Remarkably, diffuse localization of GEF-H1 was also observed upon knockdown of KANK1 in THP1 cells (Figure 8C) or KANK1 and KANK2 in HT1080 cells (Figure 8G and H) as well as in KANK1-depleted HUVECs (Supplementary Figure 3J), even though microtubule integrity was well preserved. Moreover, overexpression of KANK1-KN talin-binding domain reproduced the effect of KANK1 knockdown on GEF-H1 localization in THP1 cells (Figure 8D). Thus, not only the disruption of microtubules but also the uncoupling of microtubules from integrin adhesions led to release of GEF-H1 from microtubules. In line with previous data ^47,50^, we hypothesize that an increase of RhoA-GTP levels in KANK knockdown cells (Supplementary Figure 8) occurs due to release from microtubules and subsequent activation of GEF-H1. Thus, we investigated whether GEF-H1 depletion prevents the effects of KANK knockdowns on the myosin-II filaments and integrin adhesions in both THP1 and HT1080 cells.

GEF-H1 knockdown THP1 cells (Figure 9A) essentially preserved the phenotype of control cells, having myosin-II filaments at the periphery and podosomes occupying the more central area of the cells (Figure 9B). The integrity of the microtubule network was also unchanged (Supplementary Figure 11A). Nocodazole-induced disruption of microtubules in GEF-H1 knockdown cells was as efficient as in control cells (Supplementary Figure 11B), but did not lead to podosome disassembly (Figure 9F and G). Accordingly, microtubule disruption in GEF-H1 knockdown cells did not result in augmentation of myosin-II filaments and focal adhesion-like structures (Figure 9C and H, Supplementary Figure 11B). Moreover, double knockdown of GEF-H1 and KANK1 (Figure 9D) also did not lead to a reduction in podosome number, unlike the knockdown of KANK1 alone (Figure 9D, F and G, Supplementary Figure 11C). Remarkably, GEF-H1 knockdown also prevented the effect of overexpression of KANK1 talin-binding domain, GFP-KANK1-KN. While GFP-KANK1-KN overexpression alone led to significant disruption of podosomes and formation of focal adhesion-like structures (Figure 1N, and Figure 9E-H), the number of podosomes in GEF-H1 knockdown cells overexpressing GFP-KANK1-KN was similar to that seen in control cells while focal adhesions were practically absent (Figure 9E-H).

In HT1080 cells lacking GEF-H1, the disruption of microtubules by nocodazole did not produce any increase of focal adhesion size (Supplementary Figure 12A-C, H). Double knockdown of KANK2 and GEF-H1 (Supplementary Figure 12D, E and H) did not increase focal adhesion size above its level in cells with knockdown of GEF-H1 alone (Supplementary Figure 12C, H). Similarly, knockdown of GEF-H1 reduced also the increase of focal adhesion size in cells overexpressing the talin-binding domain of KANK2, GFP-KANK2-KN (Supplementary Figure 12F-H).

While in control HT1080 cells, microtubule outgrowth produced transient disassembly of focal adhesions (see above Figure 5A and B), in GEF-H1-depleted HT1080 cells, focal adhesion disruption by growing microtubules was less pronounced (Supplementary Figure 12I-K, Movie S17). Thus, the depletion of GEF-H1 significantly decreased the effects of microtubules on both podosomes and focal adhesions.

To assess the reliability of the knockdown results, and exclude off-target effects, validation of the knockdown of myosin-IIA and GEF-H1 were carried out using individual siRNAs (See catalogue number in Materials and Methods) corresponding to those in the same Dharmacon SMARTpool used in our experiments throughout this study. The key experiments with individual siRNAs indeed produced the same results as were obtained using Dharmacon SMARTpool (Supplementary Figure 13). The off-target effects of Dharmacon SMARTpool KANK1 and KANK2 siRNAs were rejected by rescue experiments with full-length KANK1 and KANK2, respectively (Figure 2, Supplementary Figure 4).

## Discussion

In this study, we found a unifying mechanism for the microtubule-mediated regulation of focal adhesions and podosomes operating via KANKs- and GEF-H1-dependent local reorganization of myosin-II filaments (Figure 10). It was previously shown that microtubule disruption induced disassembly of podosomes ^35^ but augmentation of focal adhesions ^18,22^. Here, we show that these apparently opposite effects on the two integrin-mediated adhesion systems can be reproduced by uncoupling of microtubules from integrin-based adhesions via knockdown of KANK family proteins, which were shown to connect talin with protein complex trapping the microtubule plus ends ^37^. We have shown for the first time that KANK family proteins indeed are required for microtubule targeting to both focal adhesions and podosomes. Moreover, we have demonstrated that experimental manipulations affecting the KANK-dependent links between microtubules and integrin adhesions have a profound effect on the integrity and dynamics of these adhesion structures.

**Figure 10.**
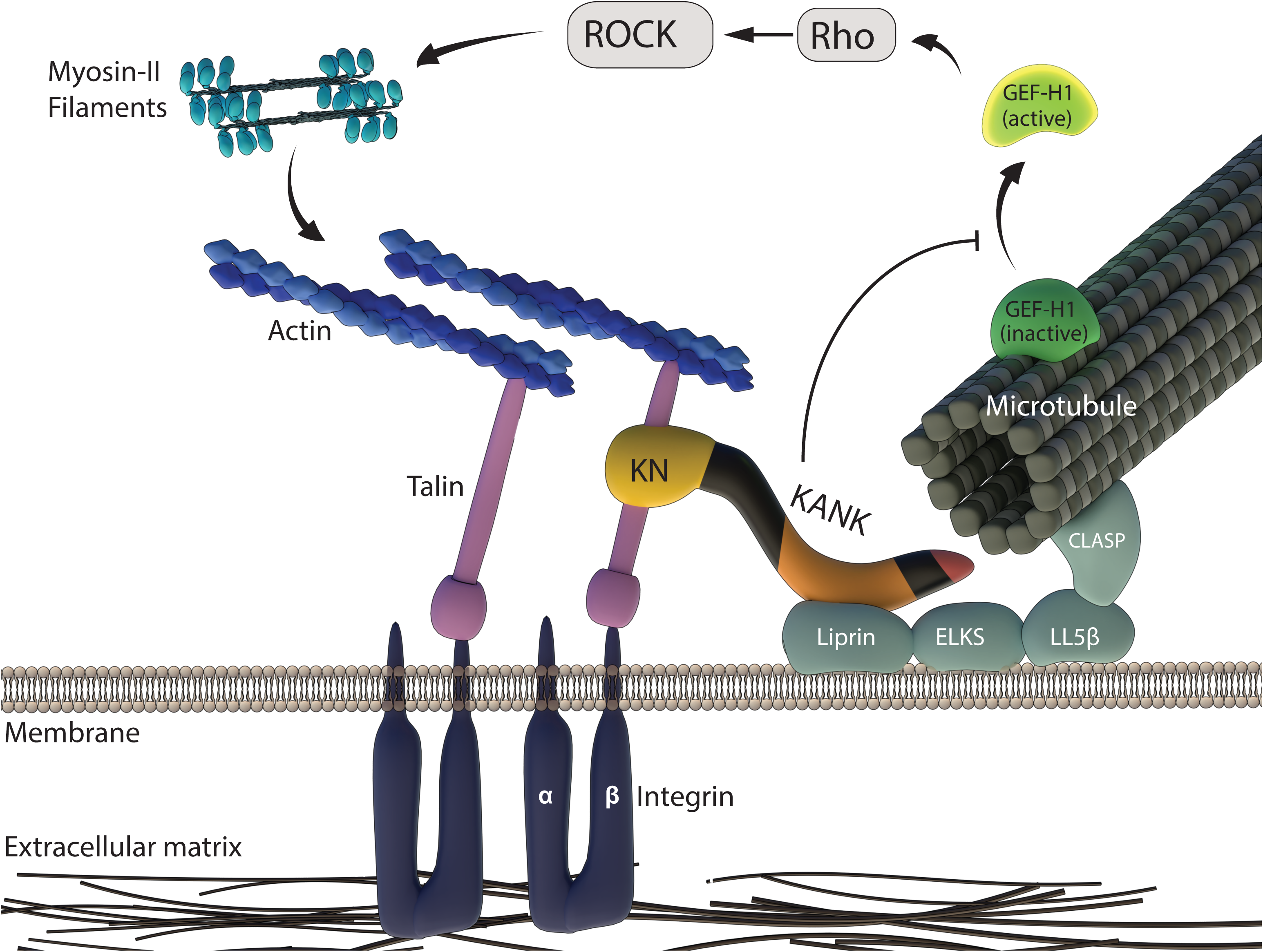
A cartoon summarizing the results of the present study. Microtubule coupled to integrin-containing adhesion structures via interaction of KANK family proteins with integrin-binding protein talin and the cortical microtubule docking complex (gray-blue) comprising of lipirins, ELKS, LL5β which captures the microtubule plus end by interacting with CLASP protein ^37^. The guanine nucleotide exchange factor GEF-H1 is associated with microtubules. Our findings suggest that coupling of microtubules to integrin adhesions via KANK suppresses GEF-H1 ability to release from microtubules. In such situations, the levels of Rho-GTP and activity of ROCK are low, and as a result, only few myosin-IIA filaments are located in the proximity of adhesions. Such conditions are permissive for podosomes but limit the growth of focal adhesions. Uncoupling of microtubule tips from integrin adhesions by depletion or displacement of KANK, leads to release of GEF-H1 from microtubules, its activation, and consequent activation of the Rho/ROCK signaling axis. As a result, the assembly of myosin-IIA filaments is significantly activated which is inimical to podosome existence. Myosin-II filaments remodel the actin cytoskeleton in the proximity of adhesions, favorable for the formation and growth of stress fiber-associated focal adhesions.

In focal adhesion-forming cells (fibroblast, epithelial and endothelial), knockdown of KANK1 or KANK2 led to augmentation of focal adhesions, mimicking the effect of total microtubule disruption. The same effect can be achieved by overexpression of talin-binding domain (KN) of either KANK1 or KANK2, which displaces endogenous KANK proteins from focal adhesions. Accordingly, we demonstrated that linking the microtubules and focal adhesions by rapamycin-induced dimerization of talin-binding domain of KANK1 with the part of KANK1 involved in microtubule trapping, led to disassembly of focal adhesions, similarly to the burst of microtubule outgrowth after nocodazole washout ^20^.

In podosome-forming cells (THP1 and RAW macrophages), the disconnection of microtubules from podosomes by overexpression of KANK1 or KANK2 talin-binding domains, led to disruption of podosomes, similarly to total microtubule disruption. Interestingly, KANK1 but not KANK2 knockdown reproduced this effect, consistently with weaker localization of KANK2 to podosomes. We found that this difference depends on the coiled-coil domain of KANK proteins known to bind liprins, the components of the membrane-associated protein complexes involved in microtubule trapping ^37^. Substitution of the coiled-coil domain of KANK2 with that of KANK1 improved the function of KANK2 as a link between microtubules and podosomes. These data show that the degree of association between KANK family proteins and liprin is an important factor regulating KANK’s function.

The screening of diverse KANK mutants revealed that the effects of KANK knockdowns can be abolished by KANK constructs that connect microtubules with talin but not by constructs that can bind either talin or microtubules alone. Altogether, these results show that uncoupling microtubules from integrin-based adhesions led to growth of focal adhesions and disruption of podosomes, faithfully reproducing the effects of total microtubule disruption. This conclusion is consistent with previous observations that knockdown of microtubule end-tracking proteins CLASP1 and CLASP2 led to stabilization of focal adhesions in fibroblasts ^23^ and destabilization of podosomes in PDBu-stimulated vascular smooth muscle cells ^51^. Indeed, it was shown that CLASPs interact with LL5β, another component of the protein complex containing liprins ^39^, which, as we have discussed above, is an important partner of KANK.

Studying the effects of microtubule disruption or uncoupling from integrin adhesions using SIM microscopy, we were able to demonstrate that the major consequence of such treatments is a dramatic increase in the amount of myosin-II filaments. Moreover, a burst of microtubule polymerization resulted in a transient decrease in the number of myosin-II filaments. We demonstrated that the effects of microtubule uncoupling or disruption on podosomes and focal adhesions can efficiently be abolished by various pharmacological and genetic manipulations affecting myosin-IIA filament assembly. In particular, pharmacological inhibition of ROCK (leading to myosin-II filament disassembly) recovered functionally active podosomes in spite of their uncoupling from microtubules or even total microtubule disruption. These observations surprisingly revealed that podosomes could degrade the matrix without immediate coupling with microtubules. It deserves further investigation then how the vesicles containing degradative enzymes (MMPs) are delivered to sites of podosome formation and function.

How does microtubule disruption or their uncoupling from adhesion structures affect myosin-II filament assembly? We investigated the role of the RhoGEF, GEF-H1 known to be associated with microtubules ^47,48,52^ in the regulation of myosin-II filament formation. We have shown that knockdown of GEF-H1 renders the level of myosin-II filament polymerization independent of microtubule integrity or their association with adhesion structures. In GEF-H1 knockdown cells, neither microtubule disruption nor KANK depletion led to podosome disassembly or focal adhesion growth. It is well recognized that GEF-H1-dependent activation of myosin-II light chain phosphorylation is mediated through activation of Rho and Rho kinase ^50^. Our data significantly extend the hypothesis of GEF-H1 regulation by microtubules ^53^, according to which GEF-H1 is inactive when associated with microtubules and undergoes activation upon microtubule disruption. Our experiments showed that GEF-H1 can be released from microtubules and apparently activated not only upon microtubule disruption but also upon uncoupling microtubules from integrin adhesions. Indeed, KANK1 or KANK2 knockdown or overexpression of KANK talin-binding domain (KN) did not induce microtubule disruption but still resulted in relocation of GEF-H1 from microtubules to cytoplasm.

Previous studies have already demonstrated that knockdown of KANK1 ^54^ and KANK2 ^55^ led to activation of Rho but overlooked the role of GEF-H1 in such activation. Our experiments suggest that not just the presence of KANK in the cytoplasm but its simultaneous link to talin and microtubules is needed for suppression of Rho activity, most probably by arresting GEF-H1 on microtubules. Thus, uncoupling of microtubules from integrin-mediated adhesions led to release and activation of GEF-H1, while targeting of the microtubules to the adhesions promotes sequestering of GEF-H1 on microtubules and consequential inactivation.

It remains to be studied which changes in biochemistry or mechanics of microtubules induced by their uncoupling from integrin adhesions are responsible for release and activation of GEF-H1. The simplest hypothesis is that contact of microtubules with focal adhesions prevents the release of GEF-H1 from microtubule tips. Another possibility is that such contact changes the organization/dynamics of the microtubule lattice preventing the release of GEF-H1 from the microtubule wall. Indeed recent studies demonstrated that microtubule bending can trigger tubulin acetylation ^56,57^, which in principle can alter the microtubule wall affinity to some associated proteins including GEF-H1.

Proteins associated with either microtubules or integrin adhesions can participate in the regulation of GEF-H1 release and association with microtubules. In particular, effects of microtubules on focal adhesions were attributed to functions of kinesin-1 ^21,58^, Eg5 kinesin ^42^, MYPT1 and HDAC6 ^59^, FAK and dynamin ^20^,^60^ and APC ^61^, while effects on podosomes are thought to involve kinesin KIF1C ^62,63^, kinesin KIF9 ^64^, and CLASPs ^51,63^. Whether and how some of these factors are regulating the process of GEF-H1 release from and sequestration by microtubules is a subject for future studies. Of note, depletion of vimentin intermediate filaments ^49^ or knockdown of EB1 ^65^ were shown to release GEF-H1 from microtubules and possibly activate it, suggesting yet other avenues for such regulation.

The question of why the myosin-II filaments affect podosomes and focal adhesions in such a different and contrasting manner also remains open. The relationship between myosin-II filaments and focal adhesions has been widely discussed in the literature ^8,13,14,16,66^. The traction force generated by myosin-II filaments promotes assembly of mechanosensitive focal adhesions ^67,68^. Indeed, in our study, the decrease of traction forces preceded the disassembly of focal adhesions induced by microtubule outgrowth, while increase of traction forces preceded the re-assembly of focal adhesion. In addition, myosin-II filaments stabilize the stress fibers associated with the focal adhesions, promoting an increase in focal adhesion size ^69,70^. The organization of the actin core of podosomes differs from the organization of actin plaques of focal adhesions. Podosome cores are filled with branched (dendritic) networks of actin filaments nucleated by WASP-Arp2/3. It is possible that such F-actin organization is incompatible with the presence of abundant myosin-II filaments undergoing rapid transformation into actomyosin bundle-like structures instead. Thus, podosomes may collapse because their dendritic networks undergo rapid transformation into actomyosin bundle-like structures. Another hypothesis is based on the fact that the existence of podosomes depends on generation of membrane curvature by podosome components interacting with the membrane such as dynamin ^71^ and FBP17 ^72^. Submembranous myosin-II filaments on the contrary appear to reduce membrane curvature ^73,74^ and therefore could antagonize podosome integrity.

The mechanism of myosin-II dependent regulation of integrin-mediated adhesions by microtubules reported in this study can contribute to the processes of cell polarization and directional migration. Microtubules at the leading edge of the cell may reduce contractility and prevent formation of large focal adhesions facilitating extension of lamellipodia ^75^. At the trailing edge of the cell, effects of microtubules on myosin-II contractility and focal adhesions could promote tail retraction ^28,76,77^. In podosome-forming cells, microtubule-driven suppression of myosin-II activity can promote formation of podosomes towards the leading edge, which may enhance directional cell movement ^78^.

Elucidation of the mechanisms underlying the effect of microtubules on integrin-mediated adhesions might be important for understanding and preventing the pathological migration of cells in the processes of metastasis. It is becoming increasingly clear that known anti-cancer effects of the drugs affecting microtubules cannot be entirely explained only in terms of inhibition of mitosis ^79,80^. There are many examples showing that such inhibitors affect either migration of tumor cells themselves or tumor-associated angiogenesis processes ^81^. The understanding that myosin-II filaments are universal effectors that mediate the microtubule-driven regulation of adhesion structures sheds a new light on the mechanism of action of these inhibitors and suggest possible new strategies for the development of druggable targets and new treatments.

In conclusion, we demonstrated here a unifying mechanism of interactions between two major elements of the cytoskeleton, microtubules and the actomyosin network, in the process of regulation of integrin-mediated adhesions. Interaction of microtubules with the cytoplasmic domains of integrin adhesions via KANK proteins regulate the release or sequestration of RhoGEF GEF-H1 which activates Rho/ROCK signaling axis that affects the formation of myosin-IIA filaments. Myosin-IIA filaments in turn operate as effectors controlling the integrin-based adhesions by generating mechanical forces that induce the growth or disassembly of these structures. This feedback regulatory loop consisting of an interwoven mix of signaling and mechanical events connecting integrin adhesions, microtubules and actomyosin provides a basis for understanding the entire cytoskeleton as a smart composite material. Detailed elucidation of the particular elements of this essential regulatory network will require further investigation.

## Acknowledgements

We thank Anna Akhmanova (Utrecht University, Netherlands) and Reinhard Fäasler (Max Plank Institute for Biochemistry, Martinsried, Germany) for KANK1 and KANK2 constructs, respectively. We are grateful to Anna Akhmanova for useful discussions and constructive criticism. This research is supported by the National Research Foundation, Prime Minister’s Office, Singapore and the Ministry of Education under the Research Centres of Excellence programme (A.D.B, N.B.M.R., T.V., and N.M.) and Singapore Ministry of Education Academic Research Fund Tier 3 (A.D.B, Y.N.) MOE Grant No. MOE2016-T3-1-002). N.B.M.R is also funded by a joint National University of Singapore-King’s College London graduate studentship. G.E.J. is supported by the Medical Research Council, UK (G1100041, MR/K015664) and the generous provision of a visiting professorship from the Mechanobiology Institute, Singapore. P.K. and Z.Z. are funded by the Ministry of Education Academic Research Fund Tier 2 (MOE-T2-1-124), the Mechanobiology Institute seed funding, the National Research Foundation Fellowship (NRF-NRFF-2011-04), and the National Research Foundation Competitive Research Programme (NRF2012NRF-CRP001-084).

## Author contributions

A.D.B. conceived and designed the project together with P.K. and G.E.J. Y.N. and N.B.M.R equally designed and performed all experiments and prepared the manuscript; Z.Z., T.V. and N.M. provided assistance in carrying out experiments and discussed results. A.D.B., G.E.J. and P.K. discussed results and prepared the manuscript.

## Materials and Methods

### Cell culture and cell transfection procedures

THP1 human monocytic leukemia cell line was obtained from Health Protection Agency Culture Collections (Porton Down, Salisbury, UK) and cultured in Roswell Park Memorial Institute media (RPMI-1640) supplemented with 10% heat-inactivated FBS and 50 μg/ml 2-Mercaptoethanol (Sigma-Aldrich) at 37°C and 5% CO_2_. The suspended THP-1 cells were differentiated into adherent macrophage-like cells with 1 ng/ml human recombinant cytokine TGFβ1 (R&D Systems) for 24 or 48 hours on fibronectin-coated glass substrates. No apparent difference between the phenotype of cells stimulated for 24 or 48 hours were detected. For the imaging samples, 35-mm ibidi (Cat. 81158) glass-bottomed dishes were coated with 1 μg/ml of fibronectin (Calbiochem, Merck Millipore) in phosphate buffered saline (PBS) for 1-2 hours at 37°C, washed with PBS twice, and immersed in complete medium prior to seeding of cells. RAW 264.7 murine macrophage cell line was purchased from American Type Culture Collection (Manassas, VA, USA) and cultured in the same media used for THP1 cells. For imaging, cells were scraped from the culture dishes, re-plated at the appropriate cell density and differentiated by addition of 100 ng/ml phorbol 12-myristate 13-acetate (PMA, Sigma-Aldrich) on glass bottom dishes for at least 24 hours.

HT1080 human fibrosarcoma cell line was obtained from American Type Culture Collection (Manassas, VA, USA) and cultured in MEM supplemented with 10% heat-inactivated FBS, Non-essential amino acid and Sodium Pyruvate (Sigma-Aldrich), in an incubator at 37°C and 5% CO_2_. HT-29 human colon adenocarcinoma cell line was gifted from Dr. Sudhakar Jha (Cancer Science Institute, Singapore) and cultured in DMEM supplemented with 10% heat-inactivated FBS and penicillin-streptomycin (Thermo Fisher Scientific), in an incubator at 37°C and 5% CO2. Primary human umbilical vein endothelial cells (HUVECs) was a gift from Dr. Roger Kamm (Singapore-MIT Alliance for Research and Technology, Singapore), which was originally purchased from American Type Culture Collection (Manassas, VA, USA). HUVECs were cultured using EGM-2 BulletKit media (catalogue no. CC-3162, Lonza, MA). All cells were plated on fibronectin-coated surfaces for 24 hours prior to imaging, or 48 hours for knockdown experiments.

Cells were transiently transfected prior to stimulation with the expression vectors plasmids using electroporation (Neon Transfection System, Life Technologies) in accordance to manufacturer’s instructions. Specifically, two pulses of 1400V of 20 ms duration were used for THP1 cells, one pulse of 950V of 50 ms was used for HT1080 cells, two pulses of 1300V of 20 ms duration for HT-29 cells, one pulse of 1200V of 40 ms duration for HUVECs and one pulses of 1680V of 20 ms duration for RAW 264.7 macrophages. For siRNA-mediated knockdown, THP1 cells were transfected at the following concentrations: 150nM for KANK1 siRNA (Dharmacon, ON-TARGETplus SMARTpool siRNA, catalogue no. L-012879-01-0005), 100nM for KANK2 siRNA (Dharmacon, ON-TARGETplus SMARTpool siRNA, catalogue no. L-027345-00-0005), 150nM for MYH9 siRNA (Dharmacon, ON-TARGETplus SMARTpool siRNA catalogue no. L-007668-00-0005), 150nM for MYH9 siRNA (Dharmacon, ON-TARGETplus siRNA catalogue no. LQ-007668-00-0002, containing following individual siRNA; J-007668-05 (#05), J-007668-06 (#06), J-007668-07 (#07) and J-007668-08 (#08)), 150nM for MYH10 siRNA (Dharmacon, ON-TARGETplus SMARTpool siRNA catalogue no. L-023017-00-0005), 150nM for GEF-H1 siRNA (Dharmacon, ON-TARGETplus SMARTpool siRNA, catalogue no. L-009883-00-0005), and 150nM for GEF-H1 individual siRNA (GEF-H1 #09 siRNA) (Dharmacon, ON-TARGETplus siRNA, catalogue no. J-009883-09-0002). For control experiments, cells were transfected with non-targeting pool siRNA (Dharmacon, ON-TARGETplus, catalogue no. D-001810-10) at a concentration similar to gene-targeted siRNAs. For siRNA-mediated knockdown in HUVECs, cells were transfected at the following concentration: 100nM for KANK1 siRNA (Dharmacon, ON-TARGETplus SMARTpool siRNA, catalogue no. L-012879-01-0005). HT1080 cells were transfected at the following concentrations: 25nM for KANK1, 25nM for KANK2 siRNA, 50nM for MYH9 siRNA and 50nM for GEF-H1 siRNA (Dharmacon, see above) using DharmaFECT 1 transfection reagent (Dharmacon, catalogue no. T-2001) following the manufacturer’s protocols.

### Plasmids

Expression vectors for fluorescent protein fusion constructs were kindly provided by several laboratories, as follows: EGFP-KANK1, EGFP-KANK1-KN, EGFP-KANK1ΔANKR and EGFP-KANK1 CC-Cter ^37^ from Dr. Anna Akhmanova (Utrecht University, Utrecht, The Netherlands); EGFP-KANK2, EGFP-KANK2-KN, EGFP-KANK2 (1-670) and EGFP-KANK2ΔKN ^28^, from Dr. Reinhard Fässler (Max Planck Institute of Biochemistry, Germany); GFP-Vinculin, mCherry-Vinculin, mTFP-vinculin, mApple-MAP4, mKO-EB3, GFP-paxillin and mApple-Paxillin from Dr. Michael W. Davidson (Florida State University, FL, USA); GFP-RhoAQ63L, from Dr. Clare M. Waterman (National Institutes of Health, USA); FLAG-ROKα1-543, from Dr. Ronen Zaidel-Bar (Mechanobiology Institute, Singapore); human GFP-Myosin regulatory light chain (MRLC), from Dr. Mark Dodding (King’s College London, UK). microtubule-binding domain of E-MAP-115/ensconsin (GFP-ensconsin) ^82^; RFP-Zyxin ^83^, mouse GFP-MRLC ^84^, mCherry-UtrCH ^85^, RFP-Lifeact and GFP-β-actin ^86^ were described previously.

For generation of EGFP-KANK2/CC-KANK1 chimeric construct, the liprin-binding coiled-coil domain corresponding to amino acid residues 188-238 of full-length KANK2 ^28^ was substituted with amino acid residues of 257-500 of full-length KANK1 ^37^. The EGFP-KANK2/CC-KANK1 construct was cloned by Epoch Life Science Inc (USA). For generation of constructs used in the rapamycin-induced dimerization of KN and ΔKN of KANK1 protein, KN domain of KANK1 corresponding to amino acid residues 1-68 ^37^ was fused with mApple fluorescence tag and FKBP12-rapamycin binding (FRB) domain of mammalian target of rapamycin (mTOR) (mApple-KN-FRB). The rest of KANK1, ΔKN, corresponding to amino acid residues 69-1352 ^37^, was fused with mEmerald fluorescence tag and FK506 binding protein (FKBP12) rapamycin-binding domain (FKBP-ΔKN-mEmerald). The mApple-KN-FRB and FKBP-ΔKN-mEmerald constructs were cloned by Epoch Life Science Inc (USA).

### Live cell observations

Pharmacological treatments were performed using the following concentrations of inhibitors or activators: 1 μM for Nocodazole (Sigma-Aldrich), 30-100 μM for Y-27632 dihydrochloride (Sigma-Aldrich), 8 μM for Thiazovivin (Sigma-Aldrich), and 1 μg/ml for Rho Activator II (CN03, Cytoskeleton), 1 μg/ml rapamycin (CAS number SB123-88-9, Santa Cruz). For THP1 cells, duration of the treatment with the inhibitors was 1 hour. In some cases, cells were pre-treated with one inhibitor for 30 min and then another inhibitor was added for additional 1 hour. For nocodazole-washout experiments, transfected HT1080 cells were plated on collagen I-coated coverslips for overnight. One hour prior to imaging, nocodazole in fresh L-15 medium (Leibovitz, Sigma-Aldrich) with 10% FBS was added to the cells. The coverslips were mounted in a perfusion chamber (CM-B25-1, Chamlide CMB chamber). Nocodazole was washed-out by fresh L-15 medium with FBS just prior to the start of the acquisition.

### Immunoblotting

Cells were lysed in RIPA buffer 48 hours after transfection and extracted proteins were separated by SDS-PAGE in 4-20% SDS-polyacrylamide gel (Thermo Fisher Scientific) and transferred to PVDF membranes (Bio-Rad) at 75V for 2 hours. Subsequently, the PVDF membranes were blocked for 1 hour with 5% non-fat milk (Bio-Rad) or bovine serum albumin (BSA, Sigma-Aldrich), then incubated overnight at 4°C with appropriate antibodies: anti-KANK1 (Bethyl Laboratories, catalogue no. A301-882A, dilution 1:1000); anti-KANK2 (Sigma-Aldrich, catalogue no. HPA015643, dilution 1:1000); Anti-non muscle myosin-IIA (Sigma-Aldrich, catalogue no. M8064, dilution 1:1000); anti-GEF-H1 (Cell Signaling Technology, catalogue no. 4145, dilution 1:1000); anti-α-tubulin (Sigma-Aldrich, catalogue no. T6199, dilution 1:3000); anti-GAPDH (Santa Cruz Biotechnology, Inc., catalogue no. sc-32233, dilution 1:3000); anti-RhoA (Santa Cruz Biotechnology, Inc., catalogue no sc-418, dilution 1:1000); anti-non muscle myosin-IIB (Developmental Studies Hybridoma Bank, CMII 23, dilution 1:1000).

Subsequently, the PVDF membranes were washed 3 times (10 minutes each) and probed by incubation for 1 hour with the appropriate secondary antibodies conjugated with horseradish peroxidase (Bio-Rad). The membranes were then washed three times (15 minutes at room temperature each), developed using Pierce™ ECL western blotting substratum (Thermo Fisher Scientific) and imaged by a ChemiDoc imaging system (Bio-Rad).

### RhoA activity assay

THP1 and HT1080 cells were lysed in RIPA buffer for five minutes on ice as described in ^87^, then centrifuged at 15,000g for five minutes at 4°C. The supernatant was incubated with Glutathione-agarose beads coated with GST-tagged Rho-binding domain of Rhotekin (provided by Dr. Boon Chuan Low (National University of Singapore) at 4°C for 30 minutes. The beads were washed three times with chilled RIPA buffer before being boiled in Laemmli buffer. Pulled-down RhoA was immunoblotted using respective antibodies as described above, and normalized to total RhoA in the whole cell lysates.

### Immunofluorescence Microscopy

HT1080 cells were pre-fixed for 3 min at 37°C using 0.3 % Glutaraldehyde and 0.2% TritonX-100 in PHEM buffer (60 mM PIPES, 27 mM HEPES, 10 mM EGTA, 8 mM MgSO_4_ × 7H_2_O, pH 7.0), and then post-fixed for 15 min at 37°C using 4 % PFA (Sigma-Aldrich) in PHEM buffer. After fixation, free aldehyde groups were quenched with 5 mg/ml Sodium borohydride (Sigma-Aldrich) for 5 min, and cells were washed 3 times for 5 min in PBS and blocked for 30 min in blocking solution (2 % Bovine Serum Albumin in PBS, Sigma-Aldrich). THP1 Cells were fixed for 15 min with 3.7% PFA in PBS, washed twice in PBS, permeabilized for 10 minutes with 0.5% triton X-100 (Sigma-Aldrich) in PBS, and then washed twice again in PBS. For microtubule visualization, cells were fixed and simultaneously permeabilized for 15 min at 37°C in a mixture of 3% PFA-PBS, 0.25% Triton-X-100 and 0.2% glutaraldehyde in PBS, and then washed twice with PBS for 10 min. Before immunostaining, samples were quenched for 15 min on ice with 1 mg/ml sodium borohydride in cytoskeleton buffer (10 mM MES, 150 mM NaCl, 5 mM EGTA, 5 mM MgCl_2_, 5 mM glucose, pH 6.1). For GEF-H1 visualization, cells were fixed with 100% methanol for 5 min at −20°C. Fixed cells were blocked with 5% BSA or 5% FBS for 1 hour at room temperature or overnight at 4°C prior to incubation with the following primary antibodies: anti-tubulin (Sigma-Aldrich, catalogue no. T6199, dilution 1:300); anti-paxillin (BD catalogue no. 610569, dilution 1:200); anti-vinculin (Sigma-Aldrich, catalogue no. V9131, dilution 1:400); anti-KANK2 (Sigma-Aldrich, catalogue no. HPA015643, dilution 1:200); anti-non muscle heavy chain of myosin-IIA (Sigma-Aldrich, catalogue no. M8064, dilution 1:800); anti-GEF-H1 (Abcam, catalogue no. ab155785, dilution 1:100); anti-non muscle myosin-IIB (Developmental Studies Hybridoma Bank, CMII 23, dilution 1:100). Samples were washed with PBS three times and incubated with Alexa Fluor-conjugated secondary antibodies (Thermo Fisher Scientific) for 1 hour at room temperature, followed by three washes in PBS. F-actin was visualized by Alexa Fluor 488 Phalloidin (Thermo Fisher Scientific), Phalloidin-TRITC (Sigma-Aldrich) or Alexa Fluor 647 Phalloidin (Thermo Fisher Scientific).

### Fluorescence Microscopy

For structured illumination microscopy, two types of equipment were used: 1) spinning-disc confocal microscopy (Roper Scientific) coupled with the Live SR module ^88^, Nikon Eclipse Ti-E inverted microscope with Perfect Focus System, controlled by MetaMorph software (Molecular device) supplemented with a 100x oil 1.45 NA CFI Plan Apo Lambda oil immersion objective and sCMOS camera (Prime 95B, Photometrics), 2) Nikon N-SIM microscope, based on a Nikon Ti-E inverted microscope with Perfect Focus System controlled by Nikon NIS-Elements AR software supplemented with a 100x oil immersion objective (1.40 NA, CFI Plan-ApochromatVC) and EMCCD camera (Andor Ixon DU-897). For Total Internal Reflection Fluorescence (TIRF) microscopy, samples were imaged using a Nikon Ti-E inverted microscope controlled by Nikon NIS-Elements AR software, supplemented with a 60x 1.49 NA, Apo TIRF oil immersion objective lens and sCMOS camera (Orca Flash 4.0, Hamamatsu Photonics).

### Traction Force Microscopy

Polyacrylamide gel substrates were prepared as previously described ^89^. In brief, 25 mm round coverslips were activated by treatment with 1.2% 3-(Methacryloyloxy)propyltrimethoxysilane (Shin-Etsu Silicon, KBE-503) in 100 % methanol for 1h followed by extensive 100 % methanol washing. Then, freshly mixed solution of 0.145% bis/12% acrylamide with 100 nm yellow-green fluorescent beads (FluoSpheres, 0.1μm, Thermo Fisher Scientific, Cat# F8803) was placed on dried- and activated-coverslips to give an adhered gel with stiffness of 16kPa. Coverslips with attached gel substrate were washed three times with 0.1M HEPES-NaOH buffer (pH 7.5) and conjugated with collagen I (Thermo Fisher Scientific, A1048301) using sulfo-SANPAH (Thermo Fisher Scientific) to facilitate cell attachment. HT1080 cells were plated on the polyacrylamide gel 2 hours prior to the experiment and images of focal adhesions by RFP-Zyxin and beads were acquired by spinning disk confocal microscope equipped with a 60X water immersion objective (NA1.2, UPlanApo, Nikon) as described above. After acquisition, 0.5 % trypsin-EDTA was added into chamber to remove all cells from the substrate, and then images of beads were captured again. The traction stress field was computed as described previously ^90^.

### Micro-patterning of adhesive islands using UV-induced molecular adsorption

Adhesive islands with a diameter of 4 μm were printed in square lattices with a period of 8 μΜ. Clean glass coverslips were sealed with NOA 73 liquid adhesive to plastic-containing dishes by UV treatment for 2 minutes, and were then treated with oxygen plasma for 5 minutes. The coverslips were coated with PLL-g-PEG (PLL(20)-g[3.5]-PEG(5), SuSoS AG, Dübendorf, Switzerland) at 100 μg/mL in PBS for at least 8 hours followed by multiple washes with PBS. For micropattern printing, PRIMO system (Alveole, France) mounted on an inverted microscope (Nikon Eclipse Ti-E, Japan) equipped with a motorized scanning stage (Physik Instrumente, Germany) was used as Digital Micro-mirror Device (DMD) to create a UV pattern at 105 μm above the focal plane of the microscope. After alignment of specimen with the UV pattern by using the Leonardo software (Alveole, France), a solution of photoinitiator (PLPP, Alveole, France) was incubated on the dishes. Depending on the UV exposure and time, the UV-activated photoinitiator molecules locally cleaved the PEG chains, which permits subsequent local deposition of proteins. The dishes were washed with PBS multiple times prior to incubation with labeled fibronectin (Rhodamine Fibronectin, Cytoskeleton, Inc., 50 μg/ml) or unlabeled fibronectin mixed with Fibrinogen Alexa 647 (ThermoFisher Scientific, 50 μg/mL) at a ratio of 20:1 for 10 minutes. After several washes with PBS, HT1080 or THP1 cells were plated on these micropatterned lattices for SIM imaging.

### Matrix degradation assay

50% sulphuric-acid-washed coverslips were coated with 50 μg/ml poly-D-Lysine for 30 minutes at room temperature, and then fixed with 0.5% glutaraldehyde for 15 minutes. 0.2% gelatin warmed at 37°C was mixed with Oregon Green 488 conjugated-pig gelatin at 6:1 ratio. Coverslips were coated with gelatin mix for 10 minutes, washed with 1xPBS and then quenched with 5 mg/ml sodium borohydride for 15 minutes followed by numerous washes. For matrix degradation assay, cells were seeded on the gelatin-coated coverslips for 8 hours, fixed and stained as described above (See “immunofluorescence microscopy”). Dark spots on the fluorescently-labeled gelatin correspond to areas of matrix that were degraded by cells.

### Image processing and data analysis

The number of podosomes was quantified automatically by applying an ImageJ-based plugin for counting nuclei (ITCN) to images of the podosome core (F-actin). Quantification was validated by manual analysis of the first 5-10 cells in the specimen. The automated procedure usually detected more than 90% of the podosomes identified by manual counting. Line intensity profiles (arbitrary unit, a.u.) measuring the mean intensity of GFP or mCherry fluorescence per area (μm^2^) by ImageJ Plugin, background-subtracted and normalized per maximal intensity in the field, yielding values ranging from 0 (lowest) to 1 (highest). For the morphometric analysis of focal adhesions in HT1080 cells, a custom-written software in IDL (Harris Geospatial Inc.) was used. Immunofluorescence images of paxillin were first background-subtracted using the rolling ball algorithm. Subsequently, focal adhesion region-of -interests (ROIs) were segmented using Otsu thresholding. The software determines the areas of individual focal adhesions. The total area of focal adhesion in ROIs was calculated by summing the value of individual focal adhesions. The focal adhesions of THP1 cells were labeled with vinculin and measured using the same algorithm with RenyiEntropy or Otsu thresholding in ImageJ.

To analyze microtubules, fluorescence images of MAP4-labeled microtubules were used ^91^. The image processing was performed as described in ^92^. The enhanced images were then binarized to extract the microtubule traces by a threshold calculated using Otsu’s method. For the analysis of myosin-II filaments (labeled by GFP-MRLC) and F-actin core in podosomes (labeled by mCherry-UtrCH), segmentation was performed by binarization using Otsu’s threshold. The amounts of myosin-II, MAP4 and F-actin were computed as the total numbers of their segmented pixels.

### Gene Expression profile

The transcriptomes of THP1 and HT1080 were mapped, quantified and indicated as fragments per Kilobase million (FPKM) using the RNA-Seq technology ^93^. Briefly, the total RNA of THP1 and HT1080 cell lysates was extracted using the RNeasy Mini Kit (Qiagen) according to manufacturer’s protocol. The extracted RNA was processed and analyzed by BGI Tech Solutions Co., Ltd.

### Statistical analyses

The methods for statistical analysis and sizes of the samples (n) are specified in the results section or figure legends for all of the quantitative data. Differences were accepted as significant for *P* < 0.05. Prism version 6 (GraphPad Software) was used to plot, analyze and represent the data.

**Supplementary Figure 1.**
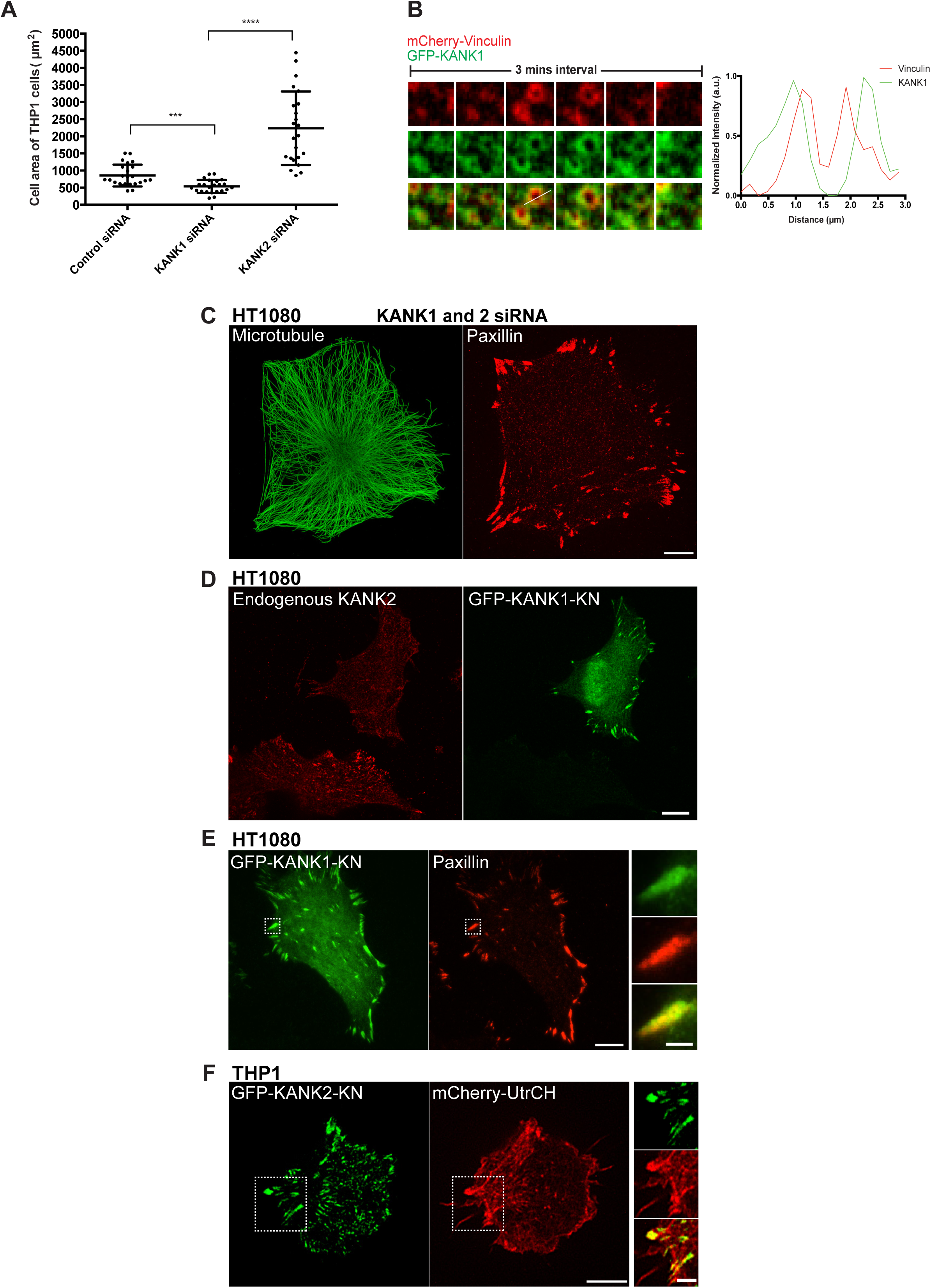
Supplementary results on KANK1 and KANK2 in THP1 and HT1080 cells. (A) Effects of knockdown of KANK1 and KANK2 on THP1 cells projected area (μm^2^). KANK1 knockdown slightly decreases cell size while KANK2 knockdown markedly increased it. Each dot corresponds to individual cell. The statistical significance of the differences was assessed using two-tailed Student’s *t*-test. (B) Localization of GFP-KANK1 to podosomes in THP1 cell. The podosomes labeled by mCherry-vinculin (red), GFP-KANK1 (green), and their merge images are shown in upper, middle and bottom rows respectively. Graph on the right represents the line scan profile of vinculin (red) and KANK1 (green) in the podosome shown in the third column. KANK1 is located to the outer ring surrounding the vinculin ring of podosome. (C) KANK1 and KANK2 double knockdown in HT1080 cell labeled with α-tubulin (green) and paxillin (red) antibodies. The cell preserved well-developed microtubule network (left) and form large focal adhesions (right). The measurements of focal adhesion area is shown in Figure 2A. (D) Overexpression of talin-binding GFP-KANK1-KN in HT1080 cell resulted in displacement of endogenous KANK2 from the focal adhesions. Two cells labeled with antibody against KANK2 (red) are shown in the left image. GFP-KANK1-KN (green) is expressed only in the upper cell (shown in the right image). Note that the lower cell which does not express GFP-KANK1-KN displays prominent KANK2-positive focal adhesions while the upper cell expressing GFP-KANK1-KN contains less of endogenous KANK2 in focal adhesions. (E) Localization of talin-binding GFP-KANK1-KN domain (green) to focal adhesions marked by mApple paxillin (red) in HT1080 cell. The enlarged images of boxed focal adhesion (on the right) demonstrate that KANK1-KN fully covers the paxillin at the focal adhesion (merged image at the bottom). Scale bars in C-E, 10 μm; magnified images in E, 2 μm. (F) Overexpression of GFP-KANK2-KN (green) in THP1 cell. The actin is labeled by mCherry-UtrCH (red). Note that GFP-KANK2-KN overexpression resulted in total disruption of podosomes and appearance of focal adhesion-like structures. The enlarged image of boxed areas on the right shows that KANK2-KN localizes to the tips of actin positive protrusions. Scale bar, 5μm.

**Supplementary Figure 2.**
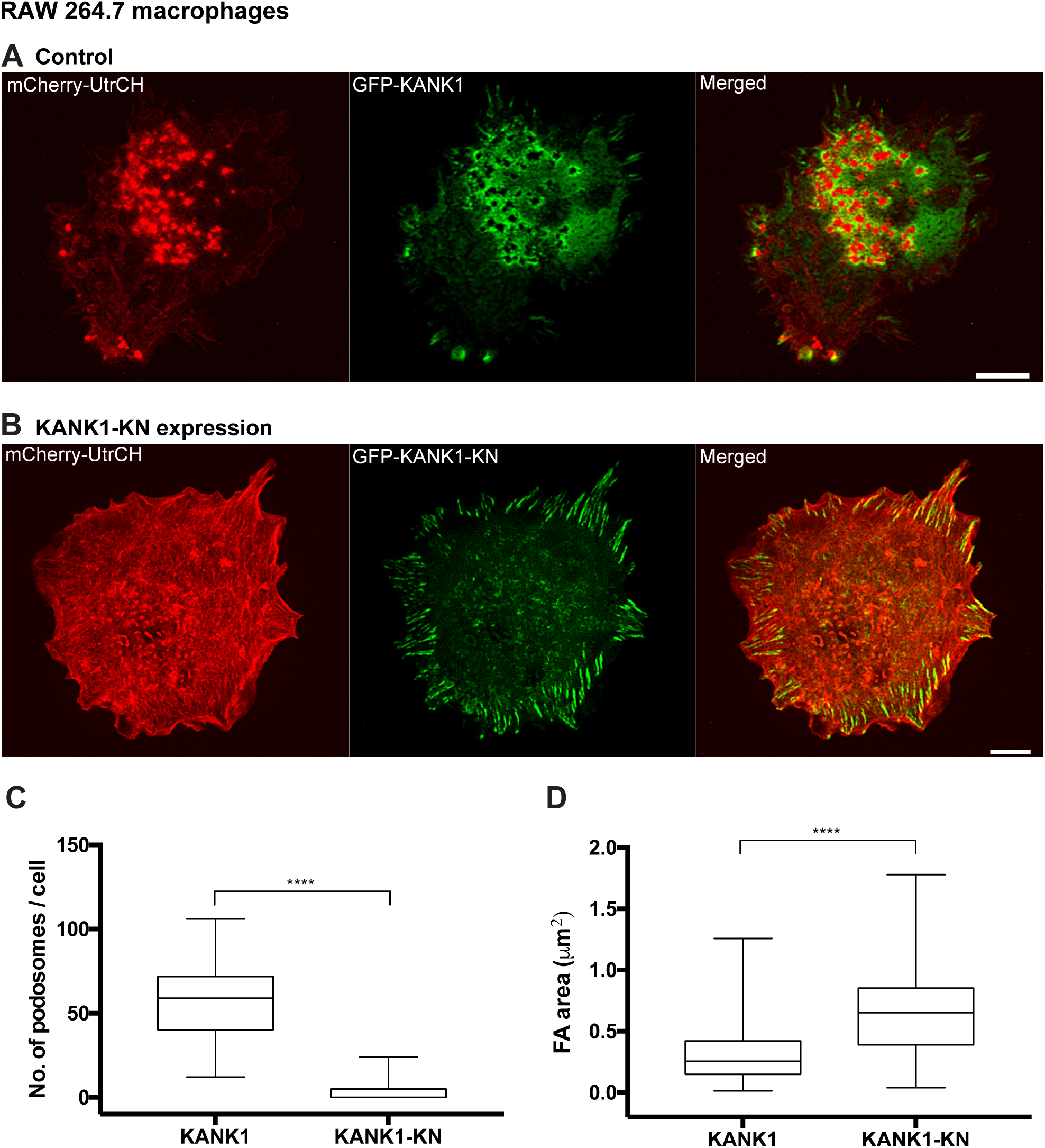
Effects of microtubule disconnection from talin in podosome-forming murine RAW 264.7 macrophages. F-actin labeled by mCherry-UtrCH (red) and GFP-KANK1 (green) in phorbol 12-myristate 13-acetate (PMA)-stimulated RAW 264.7 cell (A). Note that GFP-KANK1 localize to the adhesive rings surrounding the actin cores of podosomes. (B) PMA-stimulated RAW 264.7 cell expressing talin-binding domain of KANK1 (GFP-KANK1-KN, green) lose the podosomes and form focal adhesions instead. (C) The numbers of podosomes per cell in control and KN-expressing cells. (D) The average areas of focal adhesion-like structures in control and KANK1-KN overexpressing visualized by GFP-KANK1 or GFP-KANK1-KN, respectively. The data presented as box-and-whiskers plots and statistical significance of the difference (p-values) was calculated by two-tailed Student’s *t*-test, the range of P-values >0.05(non-significant), ≤ 0.05, ≤0.01, ≤0.001, ≤ 0.0001 are denoted by “ns”, one, two, three and four asterisks (*), respectively.

**Supplementary Figure 3.**
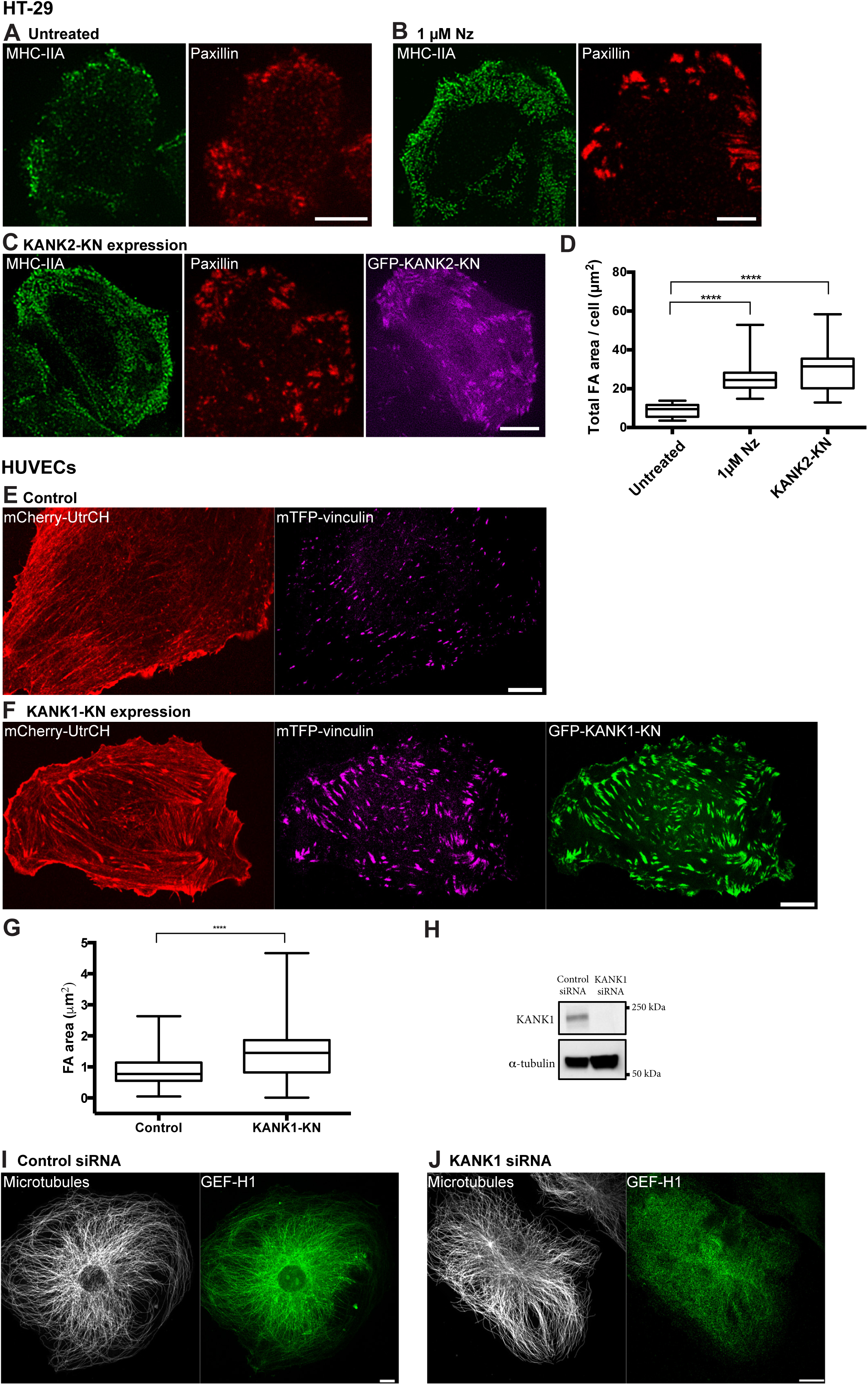
Effects of microtubule disconnection from talin in focal adhesion-forming cell types. (A-D) Human colon adenocarcinoma cell line HT-29. Myosin-IIA filaments (green) and focal adhesions labeled by paxillin (red) in untreated (A), nocodazole-treated (B), and KANK2 talin-binding domain (GFP-KANK2-KN, purple) overexpressing cells. Note the both disruption of microtubules and with nocodazole and the disconnection from integrin adhesions by GFP-KANK2-KN overexpression resulted in increased of myosin-IIA filaments number and focal adhesion size. Scale bars, 5 The quantification of total focal adhesion area per cell is shown on in (D). The myosin-II filaments and focal adhesion were visualized by immunostaining with myosin-IIA heavy chain and paxillin antibodies, respectively. (E-J) Human umbilical vein endothelial cells (HUVECs). F-actin labeled by mCherry-UtrCH (red) and focal adhesions labeled by mTFP-vinculin (purple) in control and KANK2 talin-binding domain (GFP-KANK1-KN, green) expressing cells. Note that GFP-KANK1-KN overexpression induced increase in average focal adhesion area as presented in graph (G). (H) Western blot analysis of KANK1 levels in HUVECs transfected with control and KANK1 siRNAs. α-tubulin was used as the loading control. (I and J) Microtubules (white) and GEF-H1 (green) visualized by α-tubulin and GEF-H1 antibody immunostaining, respectively, in HUVECs transfected with control siRNA (I) and KANK1 siRNA (J). Note that depletion of KANK1 resulted in the dissociation of GEF-H1 from microtubules. Scale bars, 10μm. Nz=nocodazole. The statistical significance of the difference (p-values) was estimated by two-tailed Student’s *t*-test, the range of P-values >0.05(non-significant), ≤ 0.05, ≤0.01, ≤0.001, ≤ 0.0001 are denoted by “ns”, one, two, three and four asterisks (*), respectively.

**Supplementary Figure 4.**
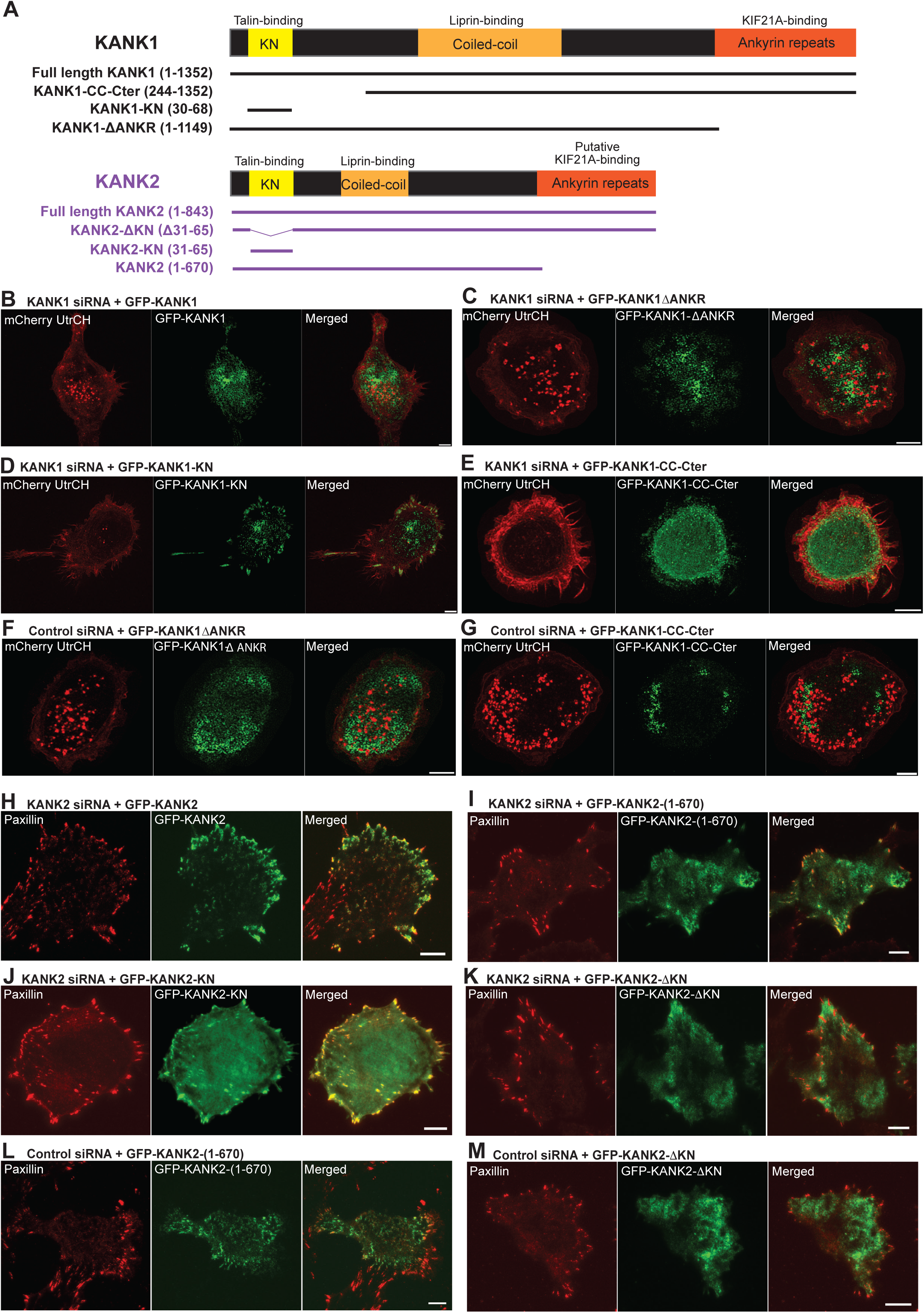
Effects of KANK1 and KANK2 constructs on podosomes and focal adhesions. (A) A map diagram depicting the domain structures of KANK1 and KANK2, and presenting their deletion mutants. KANK1 domain organization is based on data from ref ^37^. KANK1 full-length and deletion mutant GFP fusion constructs were described in ^37^. KANK2 domain structures is based on data from ref ^28^. Putative binding domain for KIF21A is indicated by analogy with KANK1. The full-length KANK2 and its deletion mutant GFP fusion constructs were described in ^28^. (B-G) Effects of full-length KANK1 and its deletion mutants on the podosomes in THP1 cells. In each panel the image of podosomes visualized by actin-labeling with mCherry-UtrCH (red) is shown on the left, localization of corresponding GFP fusion constructs of KANK1 or its deletion mutants (green) is shown in the middle, and the merged image - on the right. (B-E) KANK1 knockdown cells lacking podosomes (as shown in Figure 1G) were transfected with full-length KANK1, or KANK1 truncated constructs. (B and C) Full-length GFP-KANK1 (B) and GFP-KANK1ΔANKR (C) rescued podosomes in the KANK1-depleted cells. (D and E) GFP-KANK1-KN (D) and GFP-KANK1-CC-Cter (E) did not rescue podosomes in KANK1-depleted cells. The quantitative results corresponding to experiments shown above (B-E) are presented in Figure 2B and C. (F and G) Overexpression of GFP-KANK1ΔANKR (F) and GFP-KANK1-CC-Cter (G) in cells transfected with control siRNA did not disrupt podosomes. Scale bars, 5 μm. (H-M) Effects of full-length KANK2 and its deletion mutants on focal adhesions in HT1080 cells. In each panel the focal adhesions were visualized by paxillin antibody staining (red, left images). The localization of GFP fusion constructs of KANK2 or its deletion mutants (green) is shown in the middle images; the merged images are shown on the right. (H and I) KANK2 knockdown cells having larger focal adhesions (as shown in Figure 1E) were transfected with full-length KANK2, or KANK2 truncated constructs. Full-length GFP-KANK2 (H) and GFP-KANK2 (1-670) (I) abolished the increase of focal adhesion size. (J and K) GFP-KANK2-KN (J) and GFP-KANK2ΔKN (K) did not rescue the effect of KANK2 knockdown on focal adhesion size. See the measurements of focal adhesion sizes in Figure 2A. (L and M) The overexpression of neither GFP-KANK2 (1-670) (L) nor GFP-KANK2ΔKN (M) did not affect focal adhesion area in cells transfected with control siRNA. Scale bars, 10 μm.

**Supplementary Figure 5.**
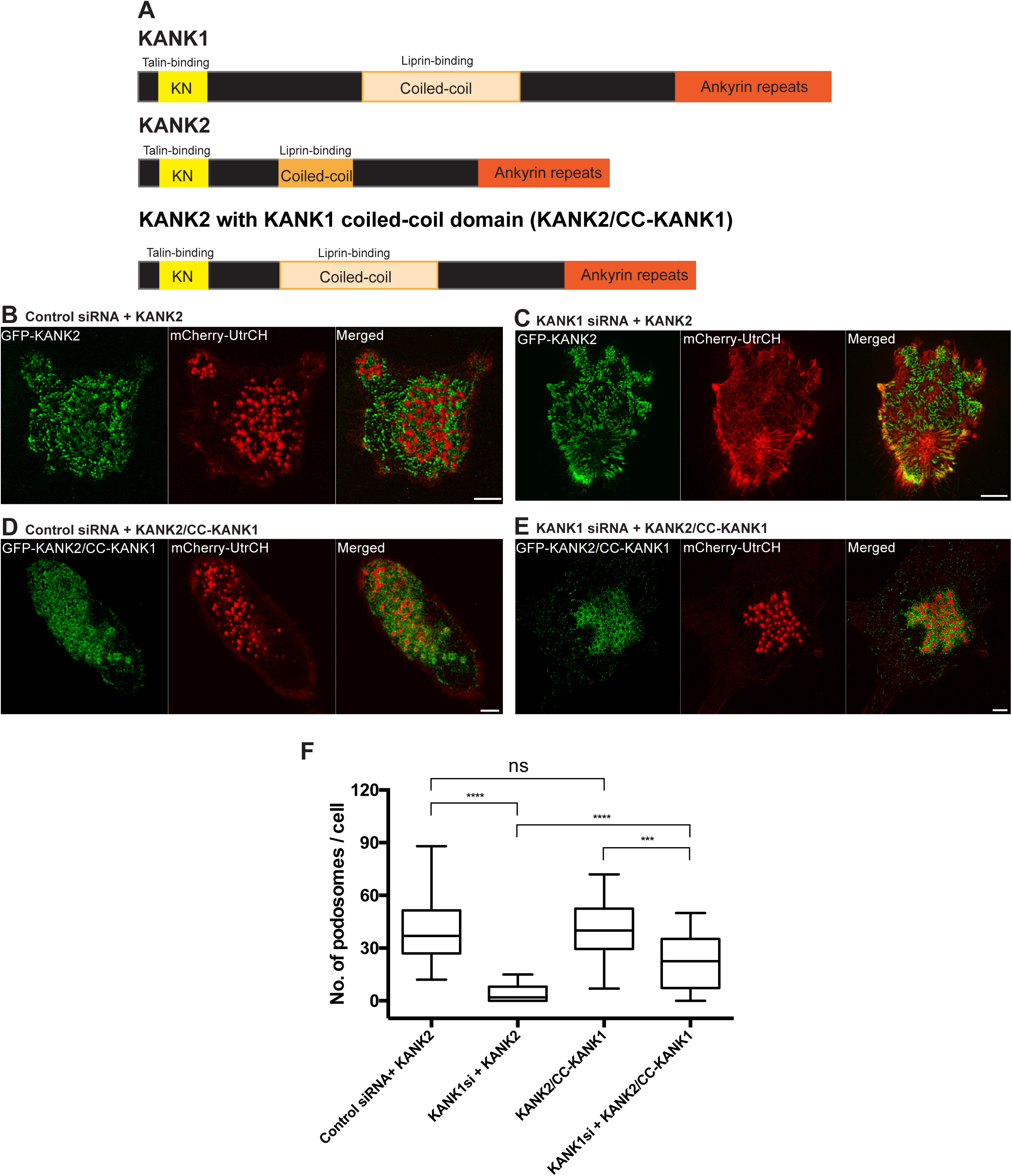
Liprin-binding domain of KANK1 is responsible for the differential effect of KANK1 and KANK2 on podosomes. (A) A map diagram depicting the domain structures of KANK1, KANK2, and a chimeric protein KANK2/CC-KANK1 designed as KANK2, in which the liprin-binding coiled-coil domain was substituted with that of KANK1. In particular, the residues 188-238 of KANK2 corresponding to its coiled-coil domain ^28^ were substituted with the residues 257-500 of KANK1 corresponding to its coiled-coil domain ^37^. (B-E) The rescue of KANK1 knockdown in THP1 cells by expression of full-length KANK2 and KANK2/CC-KANK1. In these panels the images of GFP-KANK2 or GFP-KANK2/CC-KANK1 are shown on the left (green), the images of podosomes visualized by mCherry-UtrCH (red) in the middle, while their merge images are shown on the right. (B) Expression of GFP-KANK2 in THP1 cell transfected with control siRNA did not affect podosomes. Note that GFP-KANK2 is moderately localized to podosomes. (C) Expression of GFP-KANK2 in KANK1-depleted cell did not rescue podosomes. (D) GFP-KANK2/CC-KANK1 strongly localize to podosomes in cell transfected with control siRNA. (E) Expression of GFP-KANK2/CC-KANK1 in KANK1-depleted cell rescued podosomes. The GFP-KANK2/CC-KANK1 is localized to the rescued podosomes. Scale bars, 5μm. (F) Quantification of the effect of full-length GFP-KANK2 and GFP-KANK2/CC-KANK1 on podosome number in control and KANK1 knockdown cells. Note that chimeric constructs of KANK2 with liprin-binding domain of KANK1 partially rescued podosomes in KANK1-depleted cells while full-length KANK2 did not. The data presented as box-and-whiskers plots and statistical significance of the difference (p-values) was calculated by two-tailed Student’s *t*-test, the range of P-values >0.05(non-significant), ≤ 0.05, ≤0.01, ≤0.001, ≤ 0.0001 are denoted by “ns”, one, two, three and four asterisks (*), respectively.

**Supplementary Figure 6.**
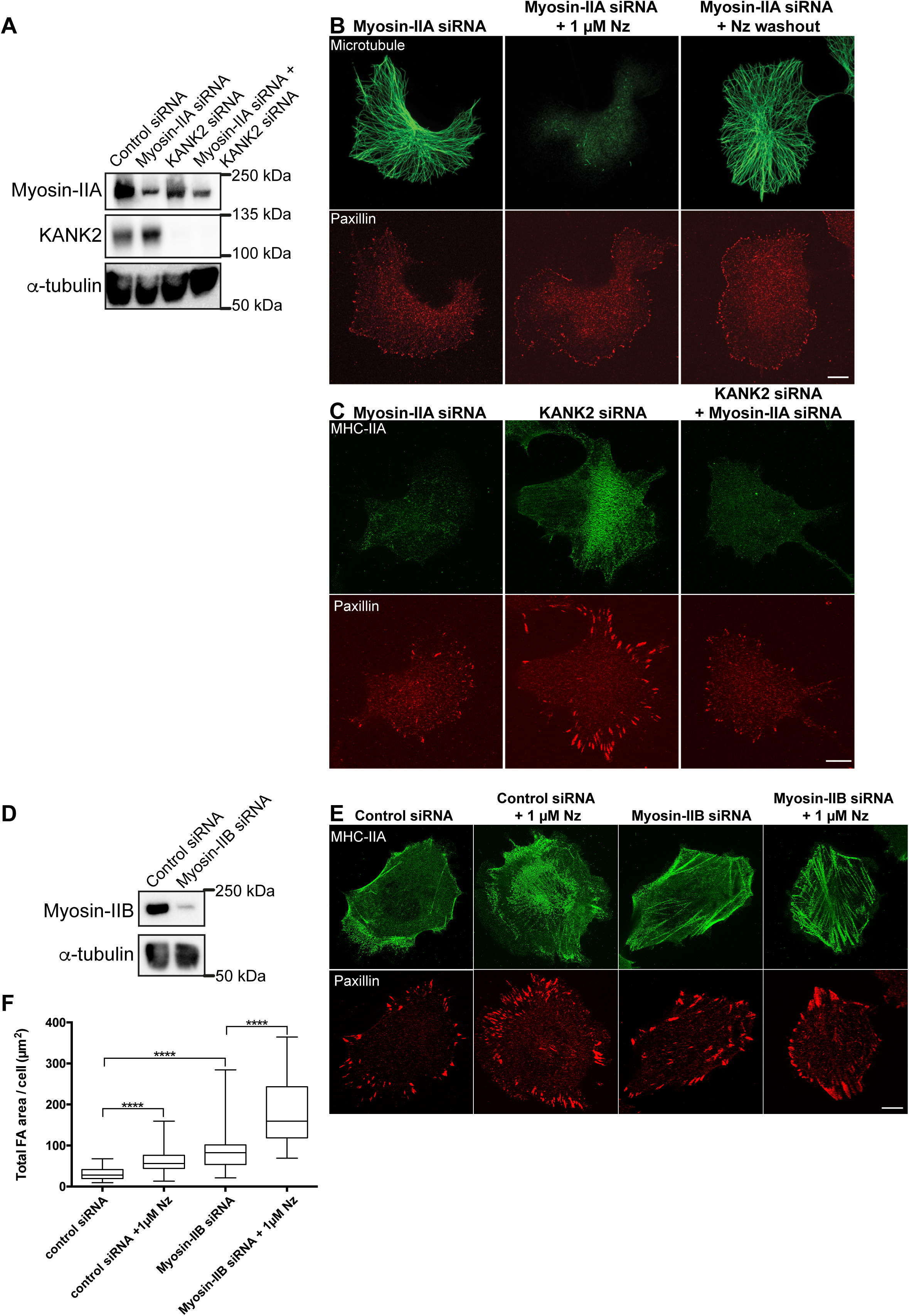
Myosin-IIA but not myosin-IIB knockdown abolished effect of microtubules on focal adhesions in HT1080 cells. (A) Western blot data showing depletion and myosin-IIA and KANK2 in HT1080 cells upon transfection with specific siRNAs. α-tubulin was used as loading control. (B) The microtubules were labeled with α-tubulin antibody staining (green, top panel) and focal adhesions in the same cells with antibody to paxillin (red, bottom panel). Untreated myosin-IIA knockdown cell is shown in the left, the similar cell treated with nocodazole in the middle and the cell after nocodazole washout on the right. Note that focal adhesion size in myosin-IIA knockdown cell is small and did not change upon microtubule disruption or outgrowth. (C) Myosin-IIA knockdown abolished the effect of KANK2 knockdown on focal adhesion size. The cells were labeled with myosin-IIA heavy chain antibody (green, upper panel) and with paxillin (red, bottom panel). Myosin-IIA knockdown cell is shown in the left pair of images, KANK2 knockdown cell in the middle, and cell with double myosin-IIA and KANK2 knockdown on the right. Notice small focal adhesion in myosin-IIA knockdown cell (left), augmented focal adhesions and myosin-II filaments in KANK2 knockdown cell (middle) and strong reduction of focal adhesion size in cell with double myosin-IIA and KANK2 knockdown (right). (D) Western blot data showing depletion of myosin-IIB in HT1080 cells upon transfection with MYH10 siRNA. α-tubulin was used as loading control. (E) Myosin-IIB knockdown did not prevent increase of focal adhesion size in nocodazole-treated cell. The cells were labeled with antibodies to myosin-IIA and paxillin as in (C). The untreated (left) and nocodazole-treated cells transfected with control siRNA (two left images) and myosin-IIB siRNA (right two images) are shown. Note that knockdown of myosin-IIB by itself somewhat increases the amount of myosin-IIA filaments and the area of focal adhesions. However, the nocodazole treatment significantly increases the total area of focal adhesions in cells transfected with either control or myosin-IIB siRNA. Scale bars in B, C and E, 10 μm. (F) Quantification of the results shown in (E). Nz=nocodazole. The statistical significance of the difference (p-values) was estimated by two-tailed Student’s *t*-test, the range of P-values >0.05(non-significant), ≤ 0.05, ≤0.01, ≤0.001, ≤ 0.0001 are denoted by “ns”, one, two, three and four asterisks (*), respectively.

**Supplementary Figure 7.**
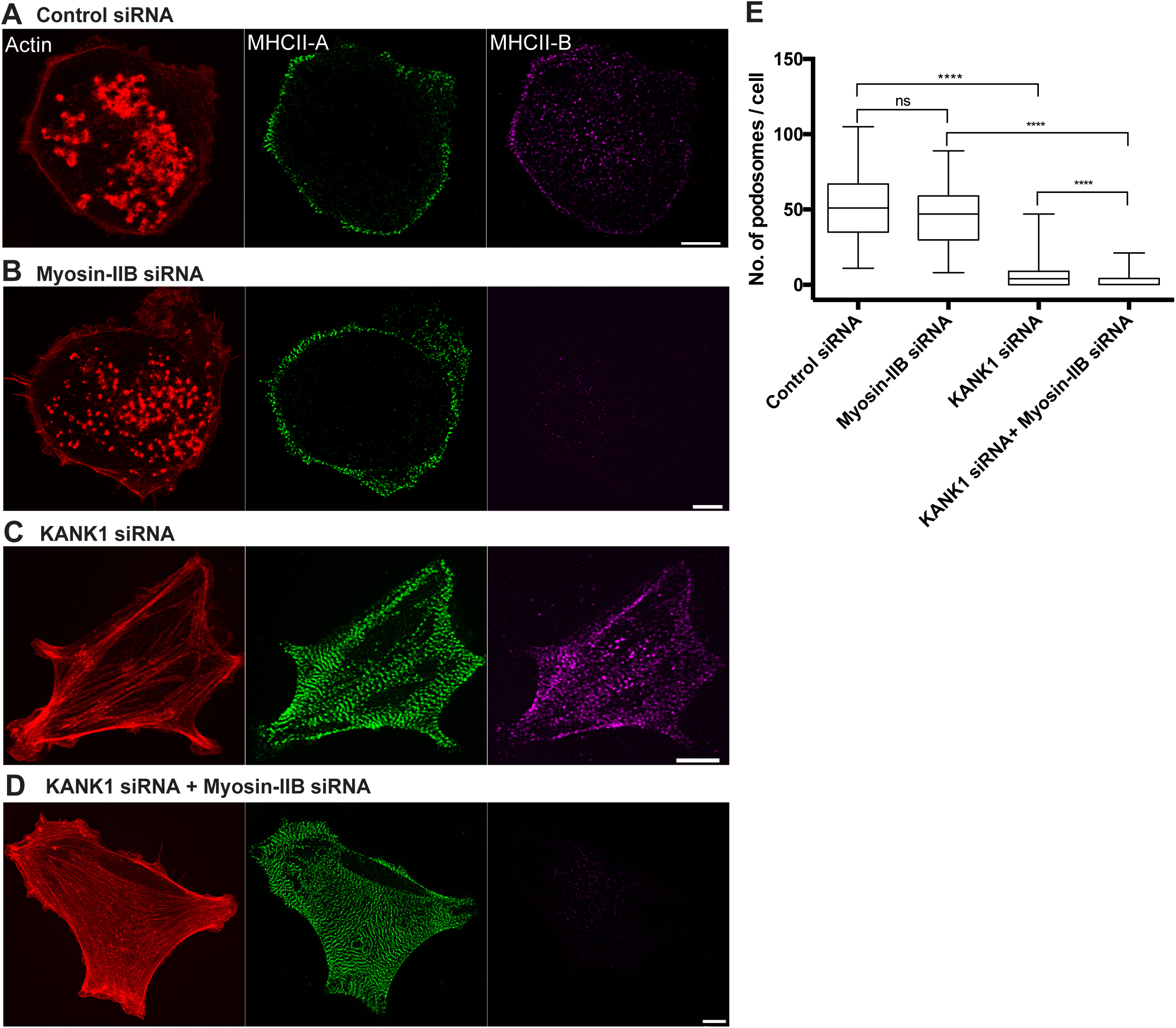
Knockdown of myosin-IIB did not prevent disruption of podosomes upon KANK1 knockdown in THP1 cells. (A-D) Podosome actin cores labeled by phalloidin (red), myosin-IIA (green) and myosin-IIB (purple) heavy chains are shown. (A) Cell transfected with control siRNA, demonstrated normal podosome distribution and peripheral localization of myosin-IIA and myosin-IIB containing filaments. Myosin-IIB staining revealed in addition some dots in the central part of the cells. (B) The knockdown myosin-IIB affected neither podosomes nor myosin-IIA-containing filaments. (C) KANK1 knockdown disrupted podosomes leading to formation of prominent actin bundles and myosin-IIA filament stacks. The distribution of myosin-IIB essentially followed that of myosin-IIA. (D) The double knockdown of KANK1 and myosin-IIB led to complete disruption of podosomes and enhancement of actin bundles and myosin-IIA filament stacks. Scale bars, 5 (E) Quantification of podosomes number in cells treated as indicated in (A-D). The data presented as box-and-whiskers plots and statistical significance of the difference (p-values) was calculated by two-tailed Student’s *t*-test, the range of P-values >0.05(non-significant), ≤ 0.05, ≤0.01, ≤0.001, ≤ 0.0001 are denoted by “ns”, one, two, three and four asterisks (*), respectively.

**Supplementary Figure 8.**
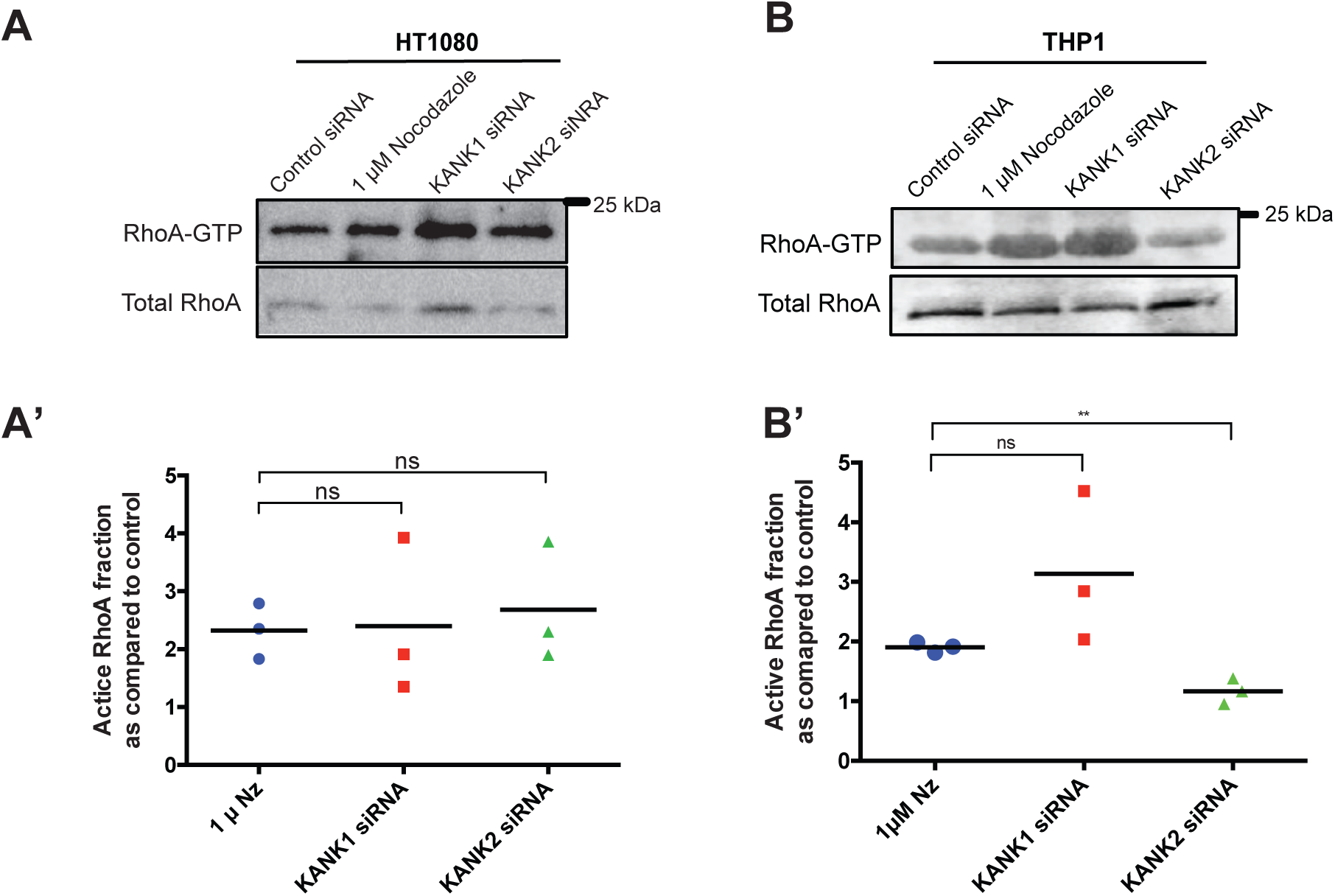
Results of pull-down assays showing effects of KANK1 and KANK2 knockdowns on RhoA activation in HT1080 and THP1 cells. Western blots of RhoA-GTP level and of total RhoA in the corresponding samples are shown in the upper and lower rows, respectively. (A and B) Effects of KANK1 and KANK2 knockdowns on the RhoA-GTP (pulled down by RBD beads) level in HT1080 (A) and THP1 cells (B) as compared to the effects of nocodazole treatment. (A) The knockdown of either KANK1 or KANK2 in HT1080 cells led to increase of RhoA-GTP level similar to nocodazole treatment as shown by quantification of the western blot data in (A’). (B) The knockdown of KANK1 but not KANK2 in THP1 cells led to increase of RhoA-GTP level similar to nocodazole treatment as shown by quantification of the western blot data in (B’). (A’ and B’) The data represent the ratios between the amounts of RhoA-GTP to total RhoA in the sample estimated as a ratio between intensities of corresponding bands. The results are then normalized to the value of such ratio in untreated cells transfected with control siRNA. Note that the average normalized values for nocodazole-treated HT1080 and THP1 cells, KANK1 and KANK2 knockdown in HT1080 cells and KANK1 knockdown in THP1 cells are more than 2-fold larger than control value (defined as unit). The statistical significance of the difference (p-values) was calculated by two-tailed Student’s *t*-test, the range of P-values >0.05(non-significant), ≤ 0.05, ≤0.01, ≤0.001, ≤ 0.0001 are denoted by “ns”, one, two, three and four asterisks (*), respectively.

**Supplementary Figure 9.**
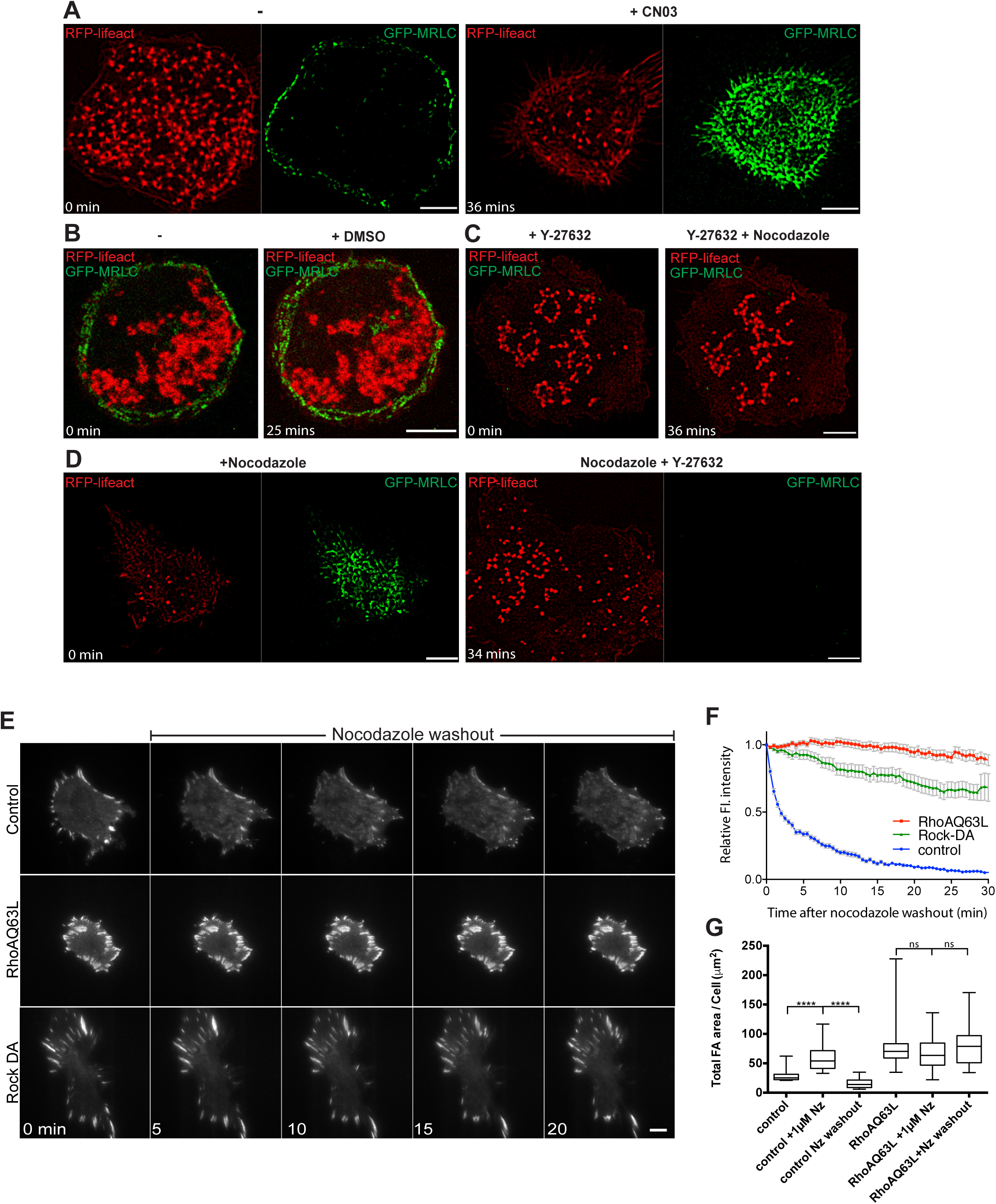
Involvement of Rho-ROCK signaling axis in microtubule-dependent regulation of podosomes and focal adhesions. (A-D) Podosomes were visualized by RFP-lifeact (red) and myosin-II filaments labeled by GFP-MRLC (green) in THP1 cell stimulated by TGFβ1. (A) Podosomes and myosin-II filaments before (left panel) and after (right panel) addition of Rho activator, CN03. Note the assembly of new myosin-II filaments and the disappearance of podosomes following CN03 addition. See Movie S10. (B) THP1 cell before (left) and after (right) 1 hour incubation with DMSO. DMSO did not change the distribution of myosin-II filaments and podosomes. See Movie S11. (C) THP1 cell pre-treated with Y-27632 for 30 minutes before (left) and after (right) 1 hour incubation with nocodazole. Pre-treatment with Y-27632 prevented burst of myosin-II filament polymerization as well as the disruptive effect of nocodazole on podosomes (cf. Figure 3A). Scale bars, 5 μm. See Movie S13. (D) THP1 cell pre-treated with nocodazole for 30 minutes before (left) and after (right) 1 hour incubation with Y-27632. Addition of Y-27632 resulted in the disassembly of myosin-II filaments (induced by nocodazole treatment), and subsequent recovery of podosomes. See Movie S14. Scale bars, 5 μΜ. (E-G) Constitutively active RhoA and ROCK1 prevented the disruption of focal adhesions following microtubule outgrowth in HT1080 cells. Scale bars, 10 μm. (E) Time-lapse sequences showing focal adhesion dynamics after nocodazole washout in control cell (top row), cell expressing constitutive active RhoA-Q63L (middle row) and cell expressing dominant active ROCK (rat Rok-alpha 1-543aa). The focal adhesions were visualized by TIRF microscopy of cells expressing mApple-paxillin. Note that microtubule outgrowth after nocodazole washout led to disassembly of the focal adhesions while constitutively active RhoA and ROCK prevented this process. See Movie S16. (F) Intensities of mApple-paxillin fluorescence in focal adhesions normalized to the intensity level in the first frame, in control, RhoA-Q63L expressing and ROCK1-DA expressing cells shown in (E). (G) Total area of focal adhesions visualized by paxillin antibody staining in control and RhoA-Q63L expressing cells. The cells were either untreated, nocodazole-treated for 1 hour, or after 30 minutes following nocodazole washout. Nz-nocodazole, NzWO-nocodazole washout. cells). n≥ 14 cells were measured for each type of treatment. The data presented as box-and-whiskers plots and statistical significance of the difference (p-values) was calculated by two-tailed Student’s *t*-test, the range of P-values >0.05(non-significant), ≤ 0.05, ≤0.01, ≤0.001, ≤ 0.0001 are denoted by “ns”, one, two, three and four asterisks (*), respectively.

**Supplementary Figure 10.**
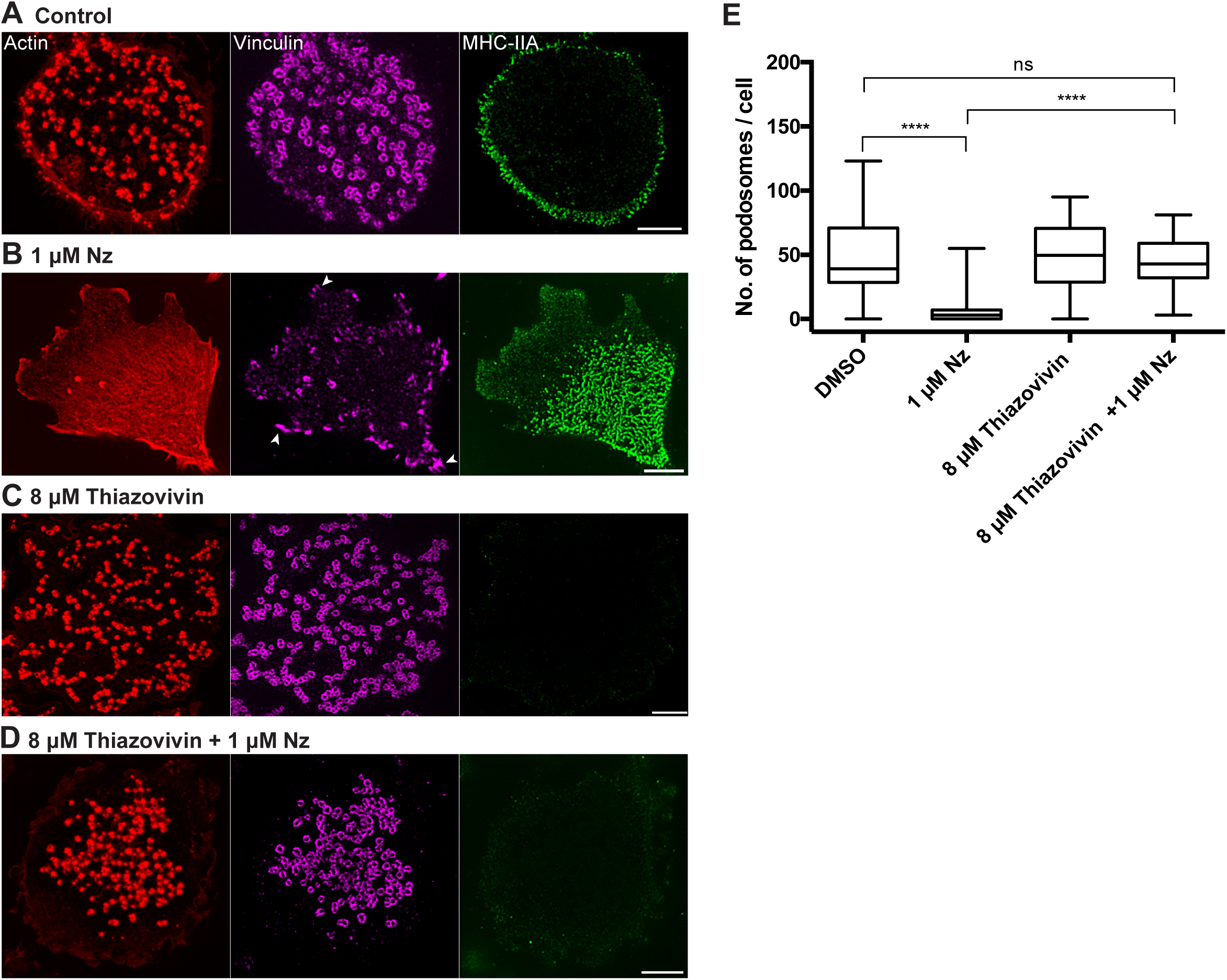
Inhibition of ROCK by thiazovivin prevents polymerization of myosin-II filaments and disruption of podosomes induced by microtubule disruption in THP1 cells. (A-D) Podosomes visualized by staining of actin cores (red) and adhesive rings (purple) and myosin-II filaments (green) in THP1 cells. (A) DMSO-treated cell (control); (B) nocodazole (Nz)-treated cell; note the disruption of podosomes and appearance of focal adhesion-like structures (arrowheads). (C) Effects of thiazovivin on podosomes and myosin-II filaments. Note that thiazovivin treatment strongly suppressed the myosin-II filaments. (D) Inhibition of ROCK by thiazovivin prevented the disruptive effect of nocodazole on podosomes. Scale bars, 5 μm. (E) Quantification of the number of podosomes per cell under the conditions indicated in A-D. The actin cores and adhesive rings of podosomes were visualized by staining of F-actin by phalloidin and vinculin by corresponding antibody. Myosin-II filaments were stained by antibody to myosin-IIA heavy chain. The data presented as box-and-whiskers plots and statistical significance of the difference (p-values) was calculated by two-tailed Student’s *t*-test, the range of P-values >0.05(non-significant), ≤ 0.05, ≤0.01, ≤0.001, ≤ 0.0001 are denoted by “ns”, one, two, three and four asterisks (*), respectively.

**Supplementary Figure 11.**
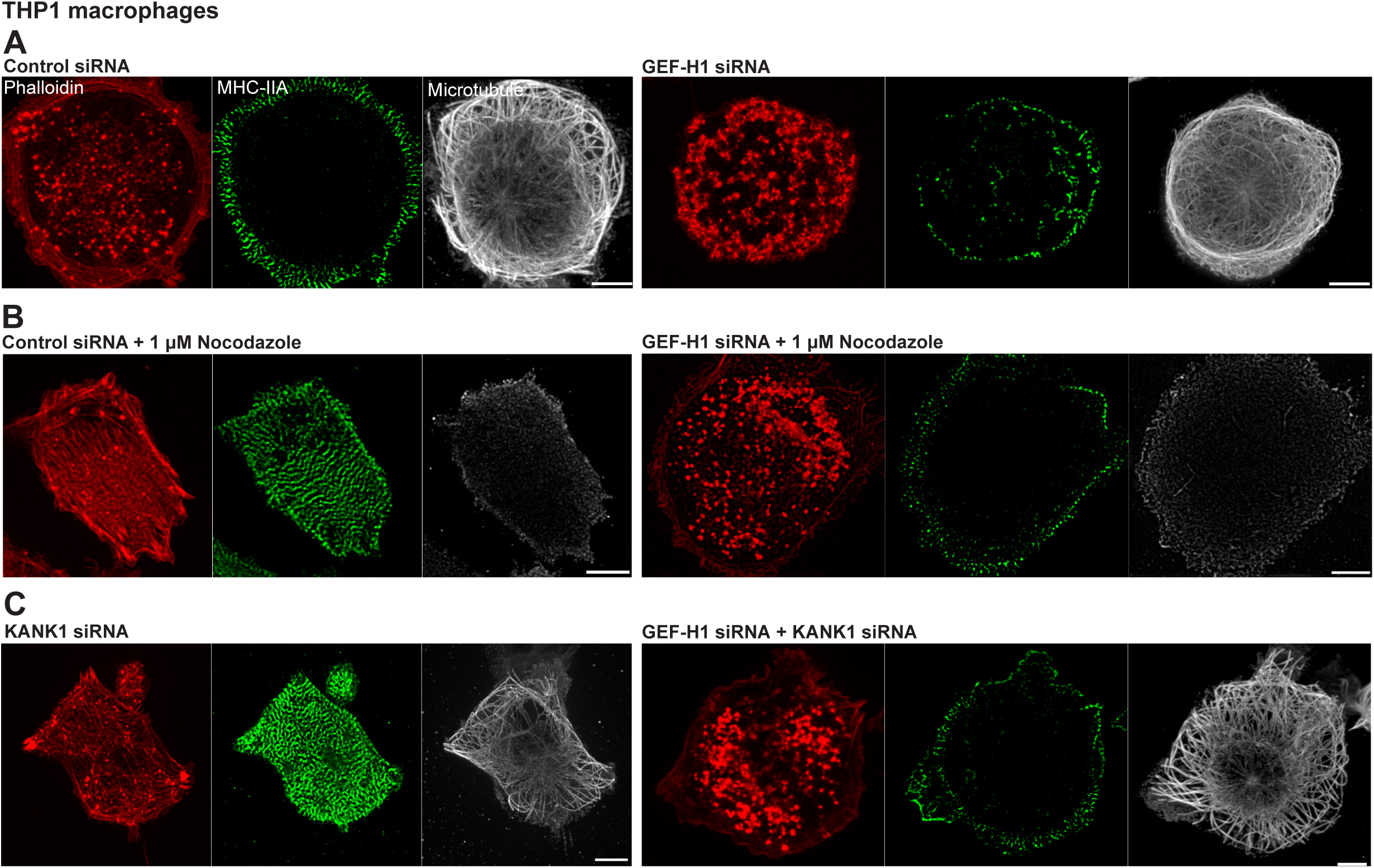
GEF-H1 knockdown did not disrupt microtubules or protect them from nocodazole but prevented the effects of nocodazole and KANK1 depletion on myosin-II filaments and podosomes in THP1 cells. (A) Control (left panel) and GEF-H1 knockdown (right panel) THP1 cells demonstrate intact podosomes visualized by the actin cores (red), peripheral rim of myosin-II filaments labeled by myosin-II heavy chain (green) and well-developed microtubule arrays (white). (B) Nocodazole treatment of control (left panel) and GEF-H1 knockdown (right panel) THP1 cells. The labeling of actin, myosin-II heavy chain and microtubules is the same as in (A). Note that nocodazole treatment disrupted podosomes and enhanced the amount of myosin-II filaments in control but not in GEF-H1 knockdown cell while disruption of microtubules was complete in both cases. (C) KANK1 knockdown THP1 cell (left panel) and cell with double knockdown of GEF-H1 and KANK1. Labeling of actin, myosin-IIA filaments and microtubules is the same as in (A) and (B). Note that KANK1 knockdown led to disassembly of podosomes and increase in the amount of myosin-II filaments. The cell lacking both GEF-H1 and KANK1 demonstrate intact podosomes and low level of myosin-II filaments. The microtubules are well preserved in both cases. Scale bars, 5 μm.

**Supplementary Figure 12.**
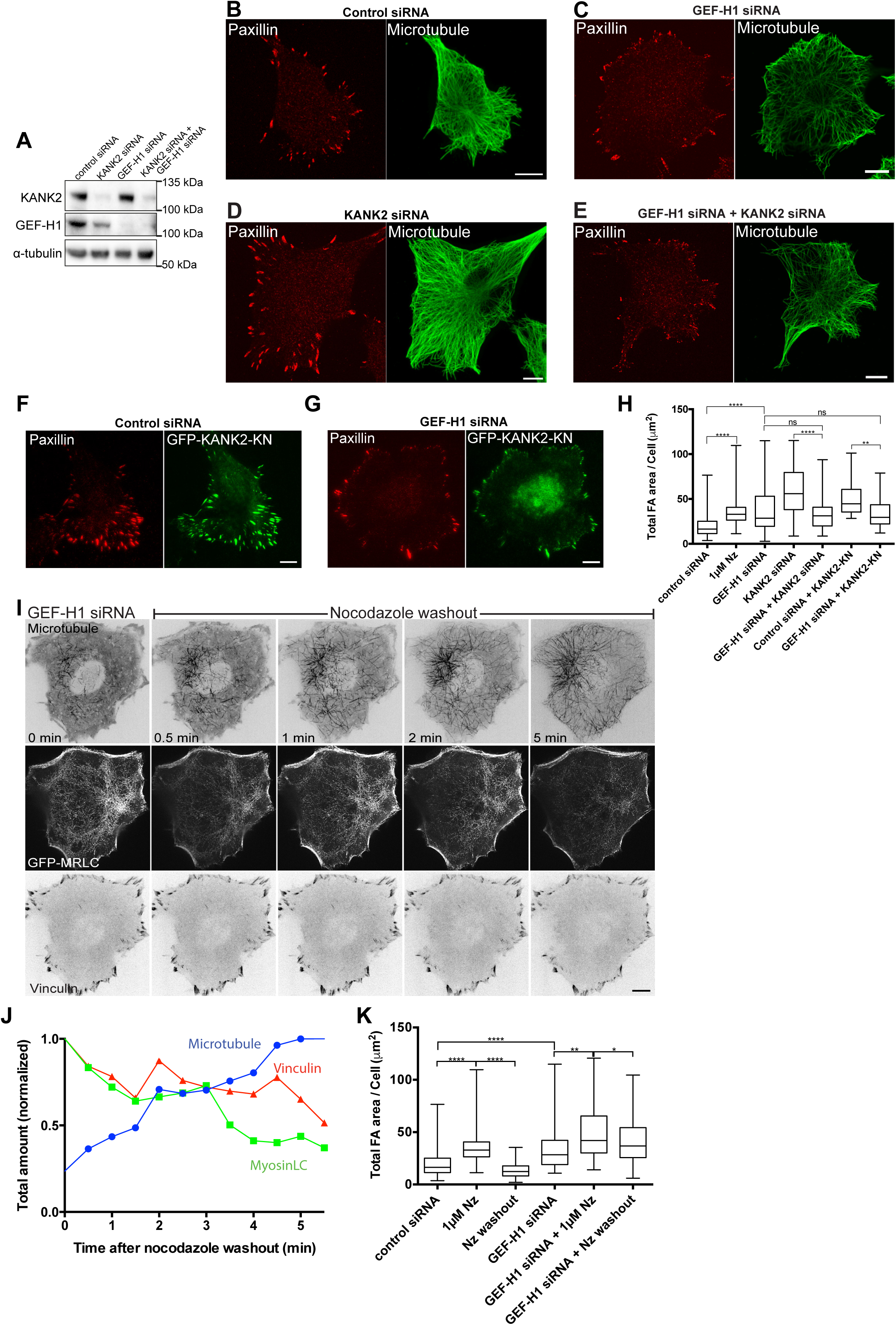
GEF-H1 knockdown prevented the effects of microtubule disconnection from focal adhesions and microtubule outgrowth on focal adhesions in HT1080 cells. (A) Western blot demonstrating siRNA-mediated depletion of GEF-H1, KANK2, and both of them in HT1080 cells. α-tubulin is used as loading control. (B-E) Focal adhesions (paxillin, red) and microtubules (α-tubulin, green) in HT1080 cells transfected with control, KANK2, GEF-H1 siRNAs, or both KANK2 and GEF-H1 siRNAs. Note that KANK2 knockdown increased the size of focal adhesions in control but not in GEF-H1 knockdown cell. The integrity of microtubules was not affected in any cases. Scale bars, 10 μm. (F and G) GEF-H1 knockdown prevented increase in focal adhesion area induced by expression of GFP-KANK2-KN. (F) Cell transfected with control siRNA and GFP-KANK2-KN (green) demonstrates large and numerous paxillin-positive focal adhesions (red). (G) The cell transfected with GEF-H1 siRNA and GFP-KANK2-KN has lower total focal adhesion area, similarly to cells transfected with GEF-H1 siRNA alone (see C). Scale bars, 10 μm. (H) Quantification of the total focal adhesion area per cell in conditions corresponding to that shown in (B-G) and in nocodazole (Nz)-treated cells. (I) The sequence of images showing the GEF-H1 knockdown cell labeled for microtubules (upper panels), myosin-II filaments (middle panels) and focal adhesions (bottom panels) in the course of microtubule outgrowth after nocodazole washout. The microtubules, myosin-II filaments and focal adhesions were labeled with mApple-MAP4, GFP-MRLC and mTFP-vinculin, respectively. See Movie S17. The images of microtubules and focal adhesions are reversed. Unlike the disruptive effect of microtubule outgrowth in control cells on myosin-II filaments and focal adhesions (Figure 4A and B), the recovery of microtubules in GEF-H1 knockdown cell was accompanied by only partial disruption of myosin-II filaments and focal adhesions (see quantification in J). (J) The microtubules (blue), myosin-II filaments (green) and focal adhesions (red) were quantified by measurements of total fluorescence intensity of cell shown in (I) at each channel. (K) The measurements of total focal adhesion area per cell in control and GEF-H1 knockdown cells upon nocodazole treatment and washout. The cells were fixed and stained with antibodies to paxillin after respective treatments. Nz=nocodazole. Not less than 45 cells were assessed for each type of treatment. The statistical significance of the difference (p-values) was estimated by two-tailed Student’s *t*-test, the range of P-values >0.05(non-significant), ≤ 0.05, ≤0.01, ≤0.001, ≤ 0.0001 are denoted by “ns”, one, two, three and four asterisks (*), respectively.

**Supplementary Figure 13.**
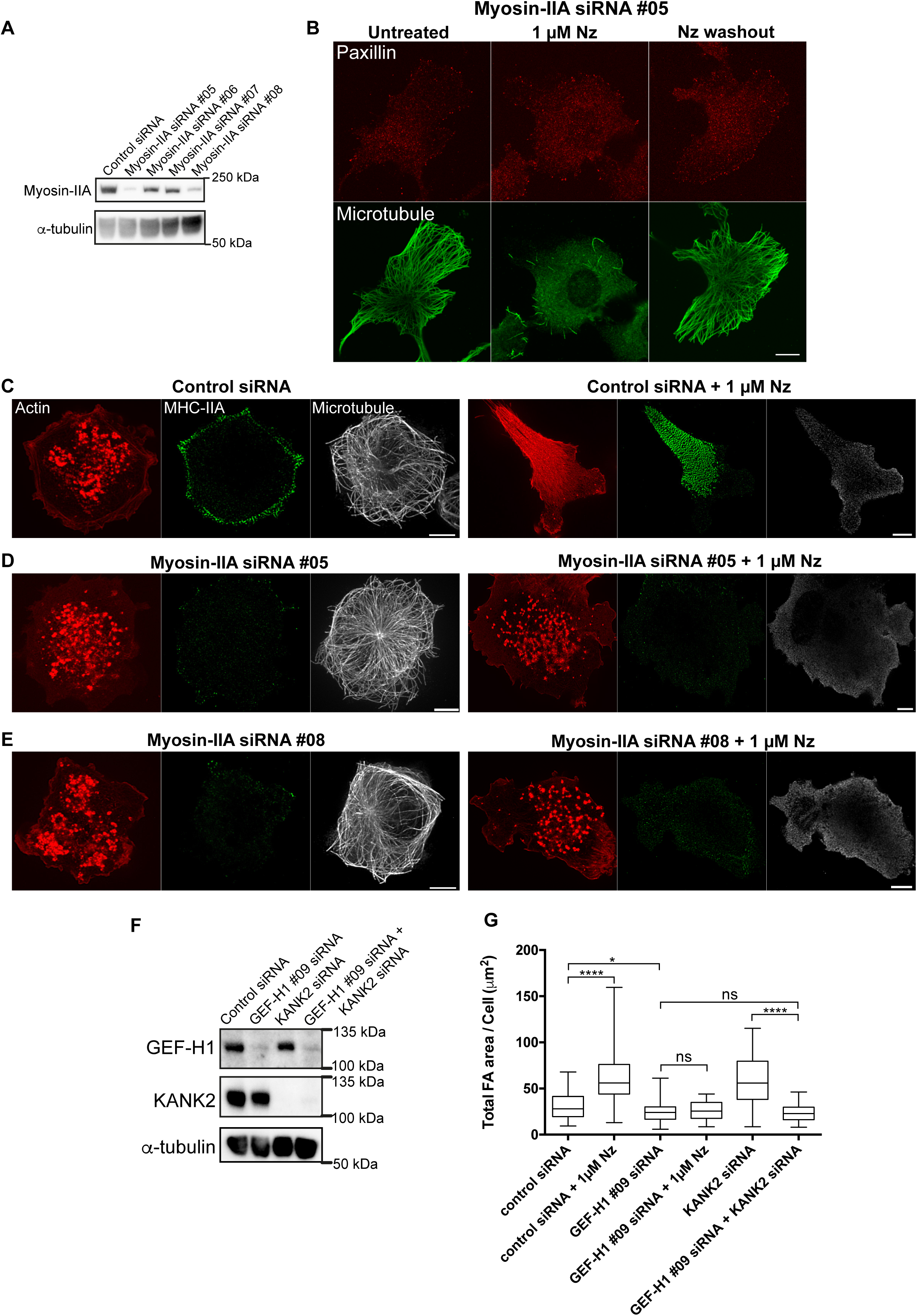
Control of off-target effects Dharmacon smartpool siRNAs against myosin-IIA and GEF-H1. (A) Western blot data showing effects of individual siRNAs on myosin-IIA heavy chain (MYH9) expression in HT1080 cells. The information corresponding to individual siRNAs (#05-#08) are shown in Materials and Methods. α-tubulin was used as loading control. (B) Cell transfection with myosin-IIA siRNA #05 prevented the effects of nocodazole and nocodazole washout on focal adhesions in HT1080 cells. Focal adhesions (red) and microtubules (green) were immunolabeled with paxillin and α-tubulin antibodies, respectively. Scale bar, 10 μm. (C-E) Myosin-II siRNAs #05 and #08 prevented the disruption of podosomes upon addition of nocodazole in THP1 cells. THP1 cell transfected with indicated siRNAs were labeled for actin (red) to visualize podosomes and immunostained for myosin-IIA heavy chain (green) and α-tubulin (white) to visualize myosin-IIA filaments and microtubules, respectively. (C) Treatment with nocodazole of cells transfected with control siRNA resulted in disappearance of podosomes and formation of numerous myosin-II filaments. (D and E) Transfection of THP1 cells with either myosin-IIA siRNA #05 (D) or #08 (E) abolished effect of nocodazole on podosomes in spite of disruption of microtubules. The effects of individual siRNAs #05 and #08 did not differ form the effect of Dharmacon smartpool presented in Figure 4 and Supplementary Figure 6. Scale bars, 5 μm. (F and G) Individual siRNA to GEF-H1 (#09) abolished the effect of nocodazole and KANK2 knockdown on focal adhesions in HT1080 cells. (F) Western blot showing the effect of individual siRNA #09 on GEF-H1 expression with and without KANK2 knockdown. α-tubulin is used as a loading control. (G) Total focal adhesion area in control, nocodazole-treated and KANK2-depleted HT1080 cells, nontransfected or transfected with GEF-H1 siRNA #09. The GEF-H1 siRNA #09 prevented the increase of focal adhesion area by nocodazole or KANK2 depletion similarly to the Dharmacon smartpool against GEF-H1 as shown in Supplementary Figure 12. The information corresponding the GEF-H1 siRNA (#09) is shown in Materials and Methods. Nz=nocodazole. The statistical significance of the difference (p-values) was estimated by two-tailed Student’s *t*-test, the range of P-values >0.05(non-significant), ≤ 0.05, ≤0.01, ≤0.001, ≤ 0.0001 are denoted by “ns”, one, two, three and four asterisks (*), respectively.

## Movie legends

**Supplementary movie 1**

The microtubule plus ends are accumulated at focal adhesion in adhesive islands of HT1080 cell transfected with control siRNA. Cell with focal adhesion labeled with GFP-vinculin (red) and microtubule plus tips labeled with mKO-EB3 (green) was imaged using SIM. Single plane close to the substrate is shown. The frames were recorded at 1-second intervals over a period of 3 minutes. Display rate is 10 frames/sec. The superimposition of frames from this movie is shown in Figure 3F (middle and right images). Scale bar, 10 μm.

**Supplementary movie 2**

The microtubule plus ends are randomly distributed over adhesive islands and spaces between them in the HT1080 cell depleted of KANK1 and KANK2. Cell with focal adhesion labeled with GFP-vinculin (red) and microtubule plus tips labeled with mKO-EB3 (green) was imaged using SIM. Single plane close to the substrate is shown. The frames were recorded at 1-second intervals over a period of 3 minutes. Display rate is 10 frames/sec. The superimposition of frames from this movie is shown in Figure 3G (middle and right images). Scale bar, 10 μΜ.

**Supplementary movie 3**

Microtubule outgrowth induces transient disassembly of focal adhesions in HT1080 cell. The imaging started immediately after nocodazole washout. The cell with microtubules labeled with GFP-ensconsin (green) and focal adhesions labeled with mApple-paxillin (red) was imaged using a spinning disk confocal microscopy. For microtubules, z-projection of all focal planes is shown; for focal adhesions, single plane near the substrate is shown. The frames were recorded at 30-seconds intervals over a period of 30 minutes. Display rate is 10 frames/sec. Scale bar, 10 μm.

**Supplementary movie 4**

Addition of 1 μg/ml rapamycin promoted the accumulation of FKBPμKN-mEmerald (red) to KN-containing focal adhesions (mApple-KN-FRB, green). The enforced dimerization by rapamycin resulted in a transient disassembly of focal adhesions marked by mTFP-vinculin (inverted black and white images). Single plane close to the substrate is shown. The frames were recorded at 30-second intervals over a period of 20 minutes using SIM. Display rate is 5 frames/sec. The movie corresponds to the time-lapse series shown in Figure 3I. Scale bar, 10 μm.

**Supplementary movie 5**

Disruption of podosomes labeled by RFP-lifeact (red, left) in THP1 cell treated with 1 μΜ nocodazole was accompanied by a massive burst of myosin-II filament assembly that aligned in stacks, as visualized by GFP-MRLC (green, right) using SIM. Single plane close to the substrate is shown. The frames were recorded at 15-second intervals over a period of 30 minutes using SIM. Display rate is 10 frames/sec. The movie corresponds to the time-lapse series shown in Figure 4A. Scale bar, 5 μm.

**Supplementary movie 6**

Myosin-II filament formation accompanies focal adhesion augmentation in HT1080 cell treated with 1 μM nocodazole. Focal adhesions are labeled with mApple-paxillin (red), myosin-II filaments are labeled with GFP-MRLC (green) and imaged using SIM. The imaging started immediately after nocodazole addition. Single plane close to the substrate is shown. The frames were recorded at 1-minute intervals over a period of 30 minutes. Display rate is 10 frames/sec. The movie corresponds to the time-lapse series shown in Figure 4B. Scale bar, 10 μm.

**Supplementary movie 7**

Microtubule outgrowth induces disassembly of myosin-II filaments in HT1080 cell. The imaging started immediately after nocodazole washout. Microtubules are labeled with mApple-MAP4 (left) and myosin II filaments are labeled with GFP-MRLC (right) and imaged using SIM. For microtubules, z-projection of all focal planes is shown; for myosin-II filaments, single plane near the substrate is shown. The frames were recorded at 15 seconds intervals over a period of 10 minutes. Display rate is 10 frames/sec. The movie corresponds to the time-lapse series shown in Figure 5A. Scale bar, 10 μm.

**Supplementary movie 8**

Changes in traction forces after microtubule outgrowth in HT1080 cell. The imaging started immediately after nocodazole washout. Traction stresses (left) computed as described in Materials and Method section are presented in spectrum scale (Pa); the scale is shown on the farmost left. Focal adhesions are labeled with RFP-Zyxin (right) and imaged using spinning disk confocal microscopy. Single plane close to the substrate is shown. The frames were recorded at 1-minute intervals over a period of 31 minutes. Display rate is 10 frames/sec. The movie corresponds to the time-lapse series shown in Figure 5C. Scale bar, 10 μm.

**Supplementary movie 9**

KANK2 knockdown suppresses disassembly of myosin-II filaments induced by microtubule outgrowth in HT1080 cell. The imaging started immediately after nocodazole washout. Microtubules are labeled with mApple-MAP4 (left), myosin II filaments are labeled with GFP-MRLC (right) and imaged using SIM. For microtubules, z-projection of all focal planes is shown; for myosin-II filaments, single plane near the substrate is shown. The frames were recorded at 15-second intervals over a period of 5 minutes. Display rate is 10 frames/sec. The movie corresponds to the time-lapse series shown in Figure 5E. Scale bar, 10 μΜ.

**Supplementary movie 10**

Disruption of podosomes labeled by RFP-lifeact (red, left) in THP1 cell treated with 1 μg/ml CN03 was accompanied by an increased assembly of myosin-II filaments visualized by GFP-MRLC (green, right), similar to nocodazole-treated cell. Single plane close to the substrate is shown. The frames were recorded at 1-minute intervals over a period of 40 minutes using SIM. Display rate is 10 frames/sec. The movie corresponds to the time-lapse series shown in Supplementary Figure 9A. Scale bar, 5 μm.

**Supplementary movie 11**

Dynamics of podosomes labeled by RFP-lifeact (red) in THP1 cell treated with 0.2% DMSO. Note that myosin-II filaments visualized by GFP-MRLC (green) are localized to the peripheral rim of the cell and are absent at sites of podosome assemblies. Single plane close to the substrate is shown. The frames were recorded at 10-seconds intervals over a period of 30 minutes using SIM. Display rate is 10 frames/sec. The movie corresponds to the time-lapse series shown in Supplementary Figure 9B.

**Supplementary movie 12**

Inhibition of Rho kinase by addition of 30 μM Y-27632 induced disassembly of myosin-II filaments visualized by GFP-MRLC (green) but did not affect podosome dynamics in THP1 cell. Podosomes are labeled by RFP-lifeact (red) and imaged using SIM. Single plane close to the substrate is shown. The frames were recorded at 10-seconds intervals over a period of 30 minutes. Display rate is 10 frames/sec. The movie corresponds to the time-lapse series shown in Figure 6A. Scale bar, 5 μm.

**Supplementary movie 13**

Inhibition of Rho kinase by 100 μM Y-27632 prevented the nocodazole-induced burst of myosin-II filaments and disruption of podosomes in THP1 cell. The imaging started at the 30^th^ minute after addition of Y-27632 and was continued for another hour in the presence of both Y-276B2 and 1 μM nocodazole. Single plane close to the substrate is shown. The frames were recorded at 2-minute intervals over a period of 40 minutes. Podosomes and myosin II filaments are labeled with RFP-lifeact (red) and GFP-MRLC (green), respectively, and imaged by SIM. The movie corresponds to the time-lapse series shown in Supplementary Figure 9C. Scale bar, 5 μm.

**Supplementary movie 14**

Disassembly of myosin-II filaments by B0 μM Y-276B2 results in recovery of podosomes in THP1 cell pre-treated with nocodazole for B0 minutes as visualized by SIM. Podosomes and myosin II filaments are labeled with RFP-lifeact (red) and GFP-MRLC (green), respectively, and imaged by SIM. The imaging started at the B0^th^ minute after addition of nocodazole and was continued for another hour in the presence of both Y-276B2 and 1 μM nocodazole. Single plane close to the substrate is shown. The frames were recorded at 2-minute intervals over a period of 40 minutes. Display rate is 10 frames/sec. The movie corresponds to the time-lapse series shown in Supplementary Figure 9D. Scale bar, 5 μm.

**Supplementary movie 15**

Inhibition of Rho kinase by treatment with B0 μM Y-276B2 in KANK1-depleted THP1 cell induced the formation of *de novo* podosomes, approximately B0 minutes after addition of the drug. Podosomes and myosin-II filaments are labeled with RFP-lifeact (red) and GFP-MRLC (green), respectively, and imaged by SIM. Single plane close to the substrate is shown. The frames were recorded at B0-second intervals over a period of 60 minutes using SIM. Display rate is 10 frames/sec. The movie corresponds to the time-lapse series shown in Figure 6B. Scale bar, 5 μm.

**Supplementary movie 16**

Constitutively active RhoA and ROCK prevented the disruption of focal adhesions upon microtubule outgrowth in HT1080 cells. Control cell (left) and cells expressing constitutive active form of RhoA (middle) or ROCK (right) are imaged just after nocodazole washout by TIRF microscopy. Focal adhesions are labeled with mApple-paxillin. The frames were recorded at B0-second intervals over a period of B0 minutes. Display rate is 10 frames/sec. The movie corresponds to the time-lapse series shown in Supplementary Figure 9E. Scale bar, 10 μm.

**Supplementary movie 17**

GEF-H1 knockdown suppresses disassembly of myosin-II filaments and focal adhesions induced by microtubule outgrowth in HT1080. The imaging started immediately after nocodazole washout. Microtubules are labeled with mApple-MAP4 (left), myosin II filaments are labeled with GFP-MRLC (right); focal adhesions are labeled with mTFP-vinculin (right), and were imaged using SIM. For microtubules, z-projection of all focal planes is shown; for myosin-II filaments and focal adhesions, single planes near the substrate are shown. The frames were recorded at 15 seconds intervals over a period of 10 minutes. Display rate is 10 frames/sec. The movie corresponds to the time-lapse series shown in Supplementary Figure 12I. Scale bar, 10 μm.

---

**Supplementary Table 1.** Gene expression profile of HT1080 and THP1 cells presented as fragments per kilobase million (FPKM) values. See materials and method section for details of the quantification.

| Gene ID | Protein | FPKM values |  |
| --- | --- | --- | --- |
|  |  | HT1080 | THP1 |
| 23189 | KANK1 | 5.46 | 6.74 |
| 25959 | KANK2 | 18.6 | 71.97 |
| 256949 | KANK3 | 0.46 | 0.23 |
| 163782 | KANK4 | 1.37 | - |
| 23332 | CLASP1 | 8.37 | 10.23 |
| 3688 | Integrin β1 | 547.88 | 69.23 |
| 3689 | Integrin β2 | 0.61 | 217.96 |
| 7094 | Talin-1 | 98.26 | 80.24 |
| 7414 | Vinculin | 81.37 | 22.35 |
| 8496 | Liprin-β1 | 65.6 | 8.29 |
| 60 | β-actin | 3876.02 | 4328.99 |
| 999 | Cadherin-1 | 0.15 | 0.67 |

